# Rapid proteasomal degradation of mutant feline McDonough sarcoma-like tyrosine kinase-3 overcomes tyrosine kinase inhibitor resistance of acute myeloid leukemia cells

**DOI:** 10.1101/2025.06.20.660791

**Authors:** Melisa Halilovic, Mohamed Abdelsalam, Andreas O. Mieland, Sarah Neuroth, Ramy Ashry, Joanna Zabkiewicz, Michelle Lazenby, Caroline Alvares, Matthias Schmidt, Walburgis Brenner, Sara Najafi, Ina Oehme, Christoph Hieber, Yanira Zeyn, Matthias Bros, Wolfgang Sippl, Oliver H. Krämer

**Author notes:** equal first author contribution (Melisa Halilovic, Mohamed Abdelsalam). equal last author contribution (Wolfgang Sippl, Oliver H. Krämer). Correspondence and requests for materials should be addressed to Wolfgang Sippl or Oliver H. Krämer. (chemistry) or (biology). e-mail addresses of all authors.

## Abstract

**Background:** Feline McDonough sarcoma (FMS)-related receptor tyrosine kinase 3 with activating internal tandem duplications (FLT3-ITD) causes acute myeloid leukemia (AML). Targeted protein degraders for FLT3 have evolved as drugs against leukemia.

**Methods:** We synthesized and characterized MA191 as novel von Hippel-Lindau (VHL)-based proteolysis targeting chimera (PROTAC) for FLT3. We analyzed protein expression, protein degradation mechanisms, and posttranslational modifications by immunoblot. Selective proteasome modulation, an inactive stereoisomer of MA191, and siRNA confirmed the event-driven degradation of FLT3-ITD. Hematopoietic cell survival and differentiation were determined by flow cytometry using apoptosis and cell surface markers. As models, we used cultured and primary human AML cells, FLT3 inhibitor-resistant AML cells, mature blood cells, and hematopoietic stem cells. We scrutinized the databases DepMap, GEPIA2, Hemap, and HPA to assess FLT3 expression and patient survival. Experiments with *Danio rerio* larvae verified in vivo anti-leukemic activity of MA191. ANOVA and Bonferroni correction were used for statistics.

**Results:** MA191 is a rapid nanomolar apoptosis inducer in AML cells harboring FLT3-ITD (IC_50_=10.16-11.6 nM; EC_50_=0.015-0.883 µM). A stereoisomer of MA191 that cannot recruit VHL demonstrates that elimination of FLT3-ITD is superior to its inhibition. MA191 abrogates FLT3 inhibitor resistance from rebound activation of mitogen-activated kinases. Rapid depletion of FLT3-ITD by MA191 (DC_50_=10 nM) requires VHL, neddylation, and the pro-apoptotic BH3-only protein BIM. Reduction of FLT3-ITD by MA191 precedes apoptosis. This reveals an apoptosis-independent function of BIM on protein stability. Leukemia cells express more FLT3 than healthy cells (n=3675/n=1249) and FLT3 expression is associated with worse AML patient survival (p=0.0099). MA191 does not harm blood cells and bone marrow progenitor cells and does not disturb myeloid blood cell differentiation. In *Danio rerio*, MA191 halts AML cell proliferation without significant toxicity. Anti-leukemic effects of MA191 are not susceptible to anti-apoptotic effects of human stromal cells and mutations in the tyrosine kinase of FLT3-ITD that confer resistance to selective FLT3 inhibitors.

**Conclusions:** these insights and the disclosure of the structure of MA191 provide a framework for an improved design of PROTACs that target mutant FLT3 and are not vulnerable to extrinsic and intrinsic resistance mechanisms. Degradation kinetics appear as determinant of such resistance breakers.

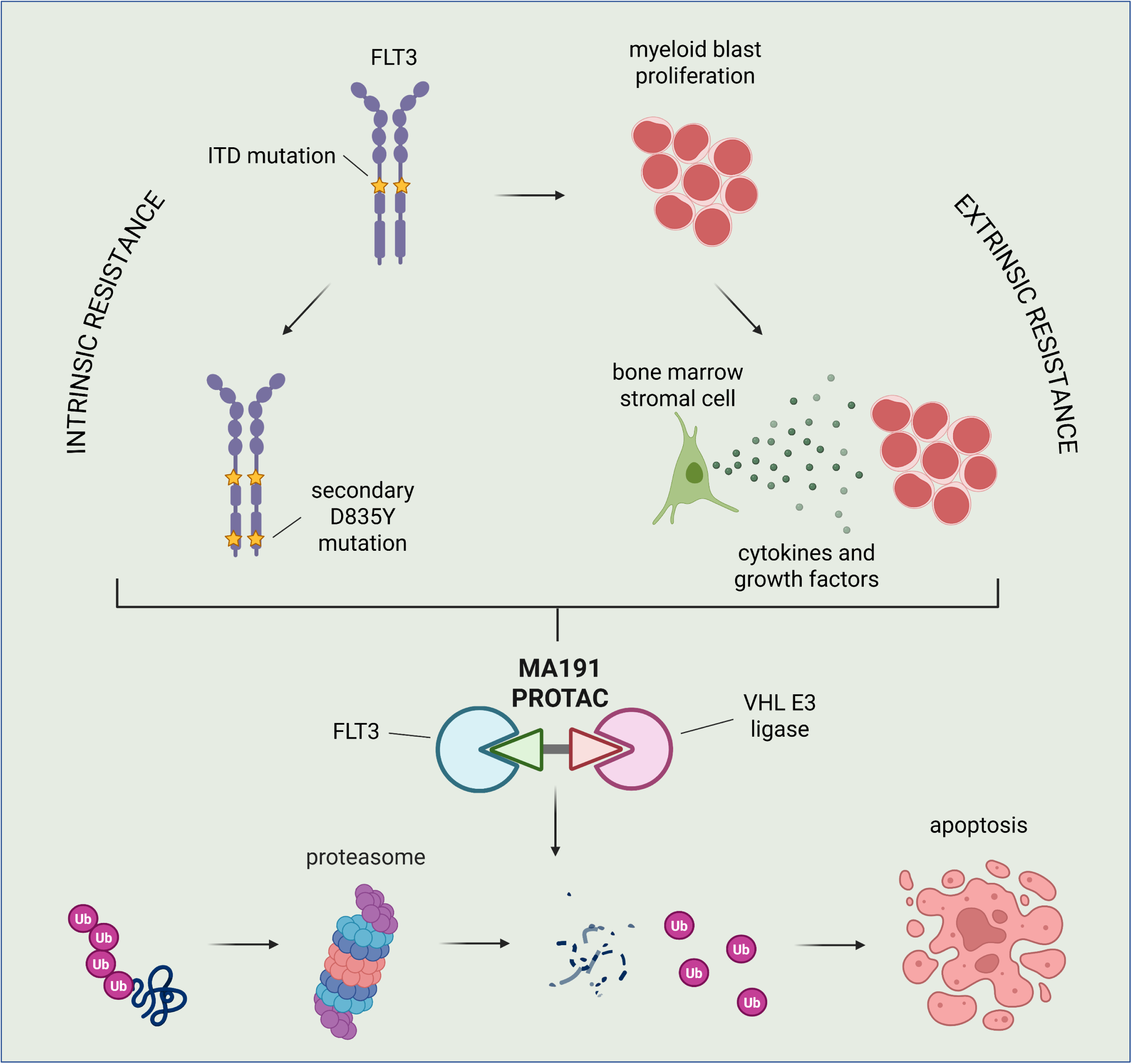

## Introduction

FMS-like tyrosine kinase 3 (FLT3) is a class III receptor tyrosine kinase (RTK) that is expressed on hematopoietic stem cells (HSCs) and early myeloid and lymphoid progenitor cells. The balanced activation of FLT3 controls the proliferation, survival, and differentiation of multipotent progenitor cells into various blood cell lineages [1]. Genetic and chromosomal changes in FLT3 can result in blocked differentiation and a rapid proliferation of hematopoietic stem and progenitor cells. Consequently, an accumulation of immature leukemic blasts in the bone marrow suppresses normal myeloid precursor cell development. This disrupts normal hematopoiesis and immune functions, and results in the sequela of leukemia [2]. Mutations in the gene encoding FLT3 account for ∼30% of acute myeloid leukemia (AML) cases. Internal tandem duplications (ITD) in the autoinhibitory juxtamembrane domain are the most prevalent FLT3 mutations. These result in the constitutive activation of FLT3 that induces leukemic cell proliferation through signal transducers and activators of transcription STAT3 and STAT5, phosphatidylinositol-3-kinase/AKT1,-2,-3, and mitogen-activated protein kinases (MAPKs). Varying ITD localization sites, peptide lengths, and sequences, as well as a high FLT3-ITD allele burden (FLT3-ITD to FLT3 wild-type ratio) create additional complexity to the disease. Such variables affect treatment responses and patient prognosis. Consequently, such variations complicate the therapeutic management of AML patients with mutant FLT3 further [3–5].

An arsenal of reversible, ATP-competitive FLT3 inhibitors has been developed for the targeted therapy of mutant FLT3-driven AML cells. Of these, midostaurin, gilteritinib, and quizartinib are clinically used. Secondary mutations in the tyrosine kinase domain (TKD) of FLT3-ITD are a cell-intrinsic mechanism of resistance that confounds the therapeutic management of patients with mutant FLT3. The most frequent secondary TKD mutation is a D835 to Y835 substitution. Substitutions at D835 occur in the activation loop of the TKD and disfavor the binding of FLT3 inhibitors, particularly type II FLT3 inhibitors (e.g., quizartinib, sorafenib). These require the inactive conformation to bind the TKD of FLT3 but have less undesired effects and are effective at lower concentrations than type I inhibitors [2, 4, 6].

The bone marrow microenvironment consists of different cell types and supplies cytokines that support hematopoiesis. Bone marrow stromal cells, including mesenchymal stem cells, osteoblasts, endothelial cells, and fibroblasts secrete cytokines and growth factors, such as interleukin-6 (IL-6), stem cell factor (SCF), CXC motif chemokine 12 (CXCL12), fibroblast growth factor 2 (FGF2), and granulocyte-macrophage colony-stimulating factor (GM-CSF). Upregulation of the CXCL12 chemokine and its corresponding receptor on leukemic cells, CXCR4, as well as the FGF2 growth factor and its receptor FGFR1, constitute an extrinsic resistance mechanism of AML cells to FLT3 inhibitors [7–10]. Leukemic cells can modify the bone marrow microenvironment to escape their elimination by immune cells. This is achieved through an inhibition of cytotoxic T cells and natural killer (NK) cells, reduced antigen-presenting major histocompatibility complex (MHC) receptor expression on dendritic cells, and an induction of immunosuppressive regulatory T cells and myeloid-derived suppressor cells [9, 11]. More work needs to be done to define how such mechanisms of leukemia cell resistance can be circumvented without damaging normal cells and tissues.

The above-summarized findings illustrate that both the bone marrow microenvironment and secondary mutations in the TKD of FLT3 contribute to AML progression and treatment failure. A further resistance mechanism that hinders the efficacy of FLT3 inhibitors relies on the therapy-induced upregulation of the FLT3 ligand, which can trigger a compensatory activation of FLT3 [12]. FLT3 inhibitor resistance can also be established through an activation of alternative signaling pathways, such as mutations in the Janus tyrosine kinase-2 (JAK2) and its downstream signaling to STAT5. One of the strategies utilized to combat these resistance mechanisms is the use of combination therapy, e.g., with CXCR4 inhibitors and a co-targeting of other kinases (e.g., JAKs, MAPKs, Aurora kinases) along with the FLT3-ITD oncogene [2, 4, 7, 8, 10, 13–15].

A new pharmacological approach to fight AML caused by FLT3-ITD is the use of FLT3 PROTACs. Such bifunctional molecules consist of a ligand for FLT3 and a ligand recruiting E3 ubiquitin ligases. These are frequently von Hippel-Lindau (VHL) protein being the substrate receptor of the cullin-2 really interesting new gene (RING)-VHL (CRL2^VHL^) E3 ligase complex or cereblon (CRBN) being the substrate receptor of the cullin-4 RING E3 ubiquitin ligase complex (CRL4^CRBN^) [16, 17]. Variable linker molecules connect the inhibitor and E3 ubiquitin ligase-recruiting moieties. Recruitment of such E3s allows PROTACs to eliminate disease-causing proteins through the cell’s proteasomal degradation machinery. This innovative mechanism offers several advantages over traditional small-molecule inhibitors, such as increased specificity and the promise to overcome resistance mechanisms through the catalytic degradation of oncoproteins [18, 19]. So far, little is known about ideal linker structures to optimize PROTACs in terms of selectivity and efficacy.

Small-molecule kinase inhibitors bind to the catalytically active sites of their target proteins. Reversible inhibitors need to continuously bind their targets to block them. In contrast, when PROTACs interact with their targets only transiently, the target protein is ubiquitinated and directed to proteasomal degradation. Upon its release, a single PROTAC molecule can induce the degradation of multiple molecules of the target protein. This recycling capacity of PROTACs allows low concentrations to achieve therapeutic effects and reduce their potential side effects [20–22]. The verification of such hypotheses requires the synthesis of molecules that structurally correspond to PROTACs except that they do not recruit the E3 ubiquitin ligase. The PROTAC-mediated, irreversible degradation of hyperactive FLT3 may prevent the emergence of secondary mutations that confer drug resistance in patients [20, 23]. An additional advantage of FLT3 PROTACs could rely on their ability to disable structural functions of FLT3-ITD. For example, it was reported that inhibited, plasma membrane-bound FLT3-ITD can confer resistance to kinase inhibitors through an activation of the MAPKs MEK and ERK [14, 15]. It has not been reported if FLT3 PROTACs can break this mechanism.

PROTACs against FLT3 are constructed from FLT3 inhibitors, such as quizartinib, gilteritinib, and dovitinib, or optimized FLT3 inhibitors that improve the formation of the ternary complex between FLT3, PROTACs, and E3 ubiquitin ligases [24–28]. Some PROTACs have higher potency or selectivity compared to their parent inhibitors. There are quizartinib-VHL, quizartinib-CRBN, and gilteritinib-CRBN-based FLT3 PROTACs. These not only differ in the kinase moieties, but also in their efficacy, the choice of E3 ligases, and linker structures [24–28]. It is often unclear if such PROTACs eliminate both FLT3-ITD and FLT3-ITD with secondary TKD mutations and if such agents affect wild-type FLT3 and normal cells.

We have reported that the molecular actions of PROTACs exceed the linear cascade of target binding and subsequent ubiquitin-dependent proteasomal degradation [29]. We found that the pro-apoptotic protein BIM was necessary for an apoptosis-associated depletion of FLT3-ITD by its PROTAC MA49. We further showed that the downregulation of protective heat shock proteins (HSPs) accompanied the PROTAC-induced depletion of FLT3-ITD in AML cells [29]. MA49 selectively kills leukemia cell lines and primary human samples that carry FLT3-ITD [29]. Here we describe MA191, an FLT3 PROTAC that is based on MA49 and contains an optimized linker. This structure increases its potency due to accelerated proteasomal degradation of mutant FLT3. MA191 shows rapid and strong activity against leukemic cell lines with FLT3-ITD and additionally acquired D835Y mutations in the FLT3-ITD allele(s). MA191 effectively induces apoptosis of AML cells that are co-cultured with human bone marrow stromal cells but does not harm the latter. MA191 is also not toxic for normal hematopoietic cells and does not impair myeloid cell differentiation. In addition, MA191 disables non-enzymatic MAPK signaling functions of FLT3-ITD and leads to a cleavage of HSP110. BIM is required for the depletion of FLT3-ITD, which precedes apoptosis induction by MA191. The in vivo efficacy of MA191 was confirmed in a *Danio rerio* early larvae model.

## Methods

### Protein lysates, immunoblot, densitometry, and antibodies

Cells were harvested on ice and centrifuged at 300xg and 4°C for 5 min. Pellets were washed with ice-cold PBS, centrifuged (300xg/5 min), and lysed in NET-N buffer (100 mM NaCl, 10 mM Tris-HCl pH 8,1 mM EDTA, 10% glycerin, 0.5% NP-40; plus complete protease inhibitor tablets (Roche, Mannheim, Germany) and phosphatase inhibitor cocktail 2 (Sigma Aldrich, Munich, Germany)) for 25 min on ice, sonicated (10 s/20% amplitude) and centrifuged (18,800xg/25 min/4°C). Protein concentrations of these whole cell lysates were measured by Bradford assay. Proteins were detected by SDS-PAGE and immunoblotting using the quantitative Odyssey Infrared Imaging System (Licor, Bad Homburg, Germany) or enhanced chemiluminescence (GE Healthcare, Freiburg, Germany) with Western Lighting Plus-ECL substrate (PerkinElmer, Waltham, USA). ImageJ was used to perform densitometric analyses.

**Table 1.** Antibodies.

| Antibody | Dilution | Catalogue number | Company |
| --- | --- | --- | --- |
| FLT3 | 1:300 | ab245116 | Abcam |
| AKT | 1:2000 | ab32505 | Abcam |
| BIM | 1:5000 | ab32158 | Abcam |
| alpha-tubulin | 1:2000 | ab176560 | Abcam |
| phospho-FLT3 (Y591) | 1:1000 | 3461 | Cell Signalling Technology |
| cleaved caspase-3 (D175) | 1:1000 | 9661 | Cell Signalling Technology |
| phospho-p44/42 MAPK (ERK1/ERK2) (T202/Y204) | 1:1000 | 9101 | Cell Signalling Technology |
| phospho-Akt (S473) | 1:1000 | 9271 | Cell Signalling Technology |
| phospho-c-Kit (Y719) | 1:1000 | 3391 | Cell Signalling Technology |
| STAT5 | 1:1000 | 33-5900 | Thermo Fisher |
| phospho-STAT5 (Y694) | 1:1000 | MA5-14973 | Thermo Fisher |
| VHL | 1:1000 | PA5-27322 | Thermo Fisher |
| HSP27 | 1:1000 | sc-13132 | Santa Cruz |
| HSP90 alpha/beta | 1:1000 | sc-13119 | Santa Cruz |
| HSP105<br>(synonym: heat shock<br>110 kDa, HSP110) | 1:1000 | sc-74550 | Santa Cruz |
| HSP70 | 1:1000 | sc-66048 | Santa Cruz |
| beta-actin | 1:1000 | sc-47778 | Santa Cruz |
| Vinculin | 1:1000 | sc-73614 | Santa Cruz |
| GAPDH | 1:2000 | sc-32233 | Santa Cruz |
| HRP-coupled anti-<br>mouse | 1:2500 | 7076 | Cell Signalling Technology |
| HRP-coupled anti-<br>rabbit | 1:2500 | 7074 | Cell Signalling Technology |
| IRDye® 680RD anti-<br>mouse | 1:10 000 | 925-68070 | Licor |
| IRDye® 680RD anti-<br>rabbit | 1:10 000 | 925-68071 | Licor |
| IRDye® 800CW anti-<br>mouse | 1:10 000 | 925-32210 | Licor |
| IRDye® 800CW anti-<br>rabbit | 1:10 000 | 925-32211 | Licor |

### Cell lines

MV4-11 (from a 10-year-old boy), MOLM-13 (from a 20-year-old male), RS4-11 (from a 32-year-old woman), and HMC-1.1/-1.2 cells (from a male mast cell leukemia patient) were gifts from Prof. Frank D. Böhmer and Dr. Sebastian Drube, University of Jena, Germany (originally from the DSMZ, Braunschweig, Germany). MOLM-13-RES and MOLM-13-RES-AC together with parental MOLM-13 cells were provided by Prof. Spiros Linardopoulos, ICR, London, UK, and are described in ref. [30]. HS-5 (CVCL_3720) are human marrow stromal cells that were immortalized by transfection with the human papillomavirus E6/E7 genes. They were a kind gift from Prof. Daniela Krause, Mainz University Medical Center, Germany. Leukemia cells were authenticated by DNA fingerprinting (DNA profiling using eight different and highly polymorphic short tandem repeats) at the Leibniz Institute (DSMZ, Braunschweig, Germany). Cells were maintained at 37°C and 5% CO_2_ in a humid atmosphere. The growth medium for MV4-11 and MOLM-13 cell lines was RPMI-1640, HMC-1.1 and HMC-1.2 cells were cultured in Iscove’s Modified Dulbecco’s Medium (IMDM), HS-5 cells were cultured in Dulbecco’s Modified Eagle Medium (DMEM). Media were supplemented with 10% fetal bovine serum (FBS) and 1% penicillin/streptomycin (Sigma Aldrich, Munich, Germany). Cell lines were confirmed to be mycoplasma-free by MycoStrip (Invivogen, Toulouse France).

### Peripheral blood mononuclear cells (PBMCs), murine bone marrow stem cells, and their differentiation into macrophages and dendritic cells

PBMCs and SCA-1/c-KIT-double positive stem cells were collected and analyzed as recently described by us [31, 32]. In brief, murine bone marrow cells were isolated from three 12-week-old male and female mice. The cells were treated with 50-100 nM MA49 or MA68. The next day, cells were harvested, washed, and stained with c-KIT, SCA-1, and a cocktail of lineage markers (CD3, CD4, CD8, NK1.1, Gr-1, CD19, CD11b). Lineage negative cells were gated for SCA-1^+^c-KIT^+^ cells and viability was assessed employing fixable viability dye 780 (FVD, negative staining indicates living cells). Additionally, bone marrow cells were seeded in 12-well plates at a density of 200,000 cells/ml and cultured for 7 days in a culture medium supplemented with either 10 ng/ml recombinant mGM-CSF to yield bone marrow-derived dendritic cells (BMDCs) or 10 ng/mL recombinant mM-CSF to differentiate bone marrow-derived macrophages (BMDMs). After plating, the cells were treated with 50-100 nM of either MA191 or MA68. The culture medium and inhibitors were replenished on days 3 and 6 of culture. After one week, cells were harvested, washed, and stained with CD86 PE (GL1, BD,#553692), CD11c PE-Cy7 (N418,eBioscience, #25-0114-81), and F4/80 FITC (BM8, eBioscience, #11-4801-85). Viability was assessed using fixable viability dye 506 (FVD-cells). After debris and doublet exclusion, total cell count was assessed and FVD-(viable) cells were further gated for CD11c or F4/80 and the mean fluorescence intensity of CD86 was assessed.

### Primary AML cells and patient characteristics

The characteristics of bone marrow cells obtained from a healthy donor and AML patients are listed in **Table 2**. Mononuclear cells were separated by Ficoll-Hypaque density gradient and subsequent dose-response assays set up with 7.5×10^5^ cells/ml cultured in IMDM medium supplemented with 10% fetal bovine serum/10% horse serum, 200 mM L-glutamine. Cells were harvested at 72 h for Cell Titer Glo cytotoxicity assays (Promega, Hampshire, UK). Luminescence was detected using a chameleon V plate reader (Hidex). Calcusyn version 2.1 (Biosoft, Cambridge, UK) and used to calculate EC_50_ responses.

**Table 2.**
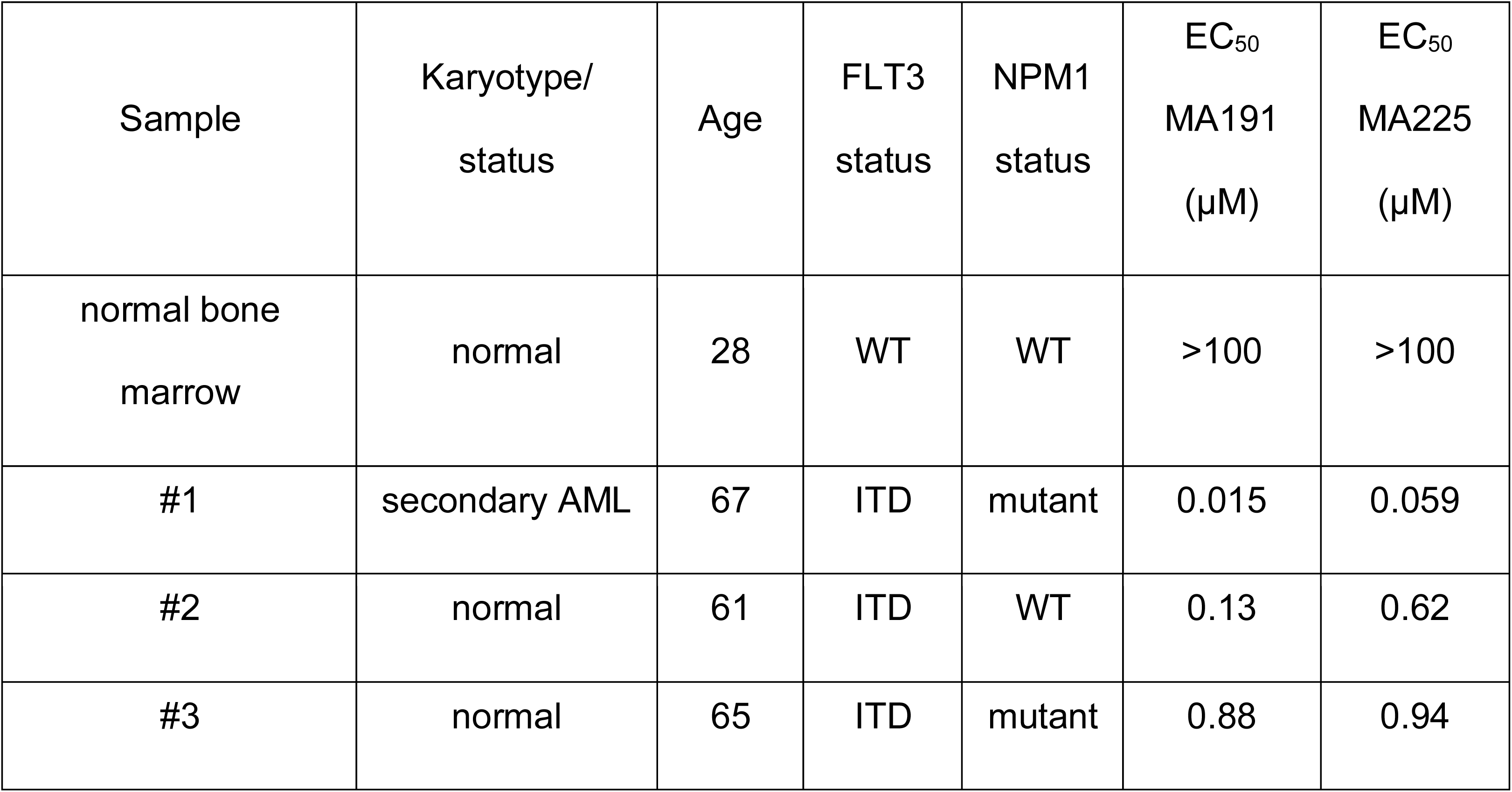
Patient age, FLT3 and NPM1 mutational status of human primary samples and EC_50_ values of MA191 and MA225.

| Sample | Karyotype/<br>status | Age | FLT3<br>status | NPM1<br>status | EC<sub>50</sub><br>MA191<br>(μM) | EC<sub>50</sub><br>MA225<br>(μM) |
| --- | --- | --- | --- | --- | --- | --- |
| normal bone<br>marrow | normal | 28 | WT | WT | >100 | >100 |
| #1 | secondary AML | 67 | ITD | mutant | 0.015 | 0.059 |
| #2 | normal | 61 | ITD | WT | 0.13 | 0.62 |
| #3 | normal | 65 | ITD | mutant | 0.88 | 0.94 |

### Inhibitors and chemicals

MA49, MA68 [29], MA191, and MA225 were synthesized by us. The analytical characterization of MA191/MA225 and their purity are described in Abdelsalam, Halilovic, Krämer, and Sippl, manuscript in preparation. Annexin-V-FITC-conjugate was from Miltenyi Biotec, Bergisch Gladbach, Germany; propidium iodide, from Sigma-Aldrich, Munich, Germany; z-VAD-FMK, Ara-C from Selleckchem, Cologne, Germany.

### Kinase binding assay

Dissociation constants (K_d_) for MA191, MA225, and MA68 at human FLT3-ITD were determined by KINOMEscan^TM^ (Eurofins DiscoverX Corporation, San Diego, CA, USA).

### Detection of apoptosis by annexin V/PI staining

Cells were collected 24-72 h after drug treatment, washed with 1x PBS, and stained with annexin V-FITC (Miltenyi Biotec, Bergisch Glabdach, Germany). After 15 min, cells were stained with PI (50 µg/ml). Flow cytometry was performed with a FACS Canto II (BD Bioscience, Heidelberg, Germany) and data were evaluated with the software tool FACSDiva 7.0 (Becton, Dickinson and Company).

### Transfections with siRNAs

MV4-11 cells were seeded in RPMI medium containing 10% FBS without antibiotics one day before electroporation. Cells were harvested and centrifuged (200xg, 5 min), the supernatant was aspirated, and cells were washed with PBS. After centrifugation (200xg, 5 min) and aspiration of PBS, cells were resuspended in 100 mL buffer R (MPK10096, Thermo Fisher, Dreieich, Germany), and siRNA solutions were added. Cells were electroporated at 1,350 V, 1 pulse for 35 milliseconds, using the Neon™ Transfection System (MPK10096, Thermo Fisher, Dreieich, Germany), and then transferred in a prewarmed culture medium. Knockdown of BIM (encoded on the *BCL2L11* mRNA) and VHL (encoded on the *VHL* mRNA) in MV4-11 cells was performed by transfecting 100 pmol SMARTpool ON-TARGETplus siRNA against *BCL2L11* (L-004383-00-0005, Dharmacon, Cambridge, UK), 100 pmol Silencer Select siRNA against *VHL* (s14789, Thermo Fisher, Dreieich, Germany), or non-targeting control siRNA (sc-37007; sc-44321, Santa Cruz, Heidelberg, Germany). After 24 h, cells were treated. Efficient knockdown was confirmed by immunoblotting.

### Zebrafish lines, embryo xenotransplantation, and treatment

The zebrafish (*Danio rerio*) embryo xenograft experiment was performed according to a previously described protocol [33], with the following specifications. MV4-11 cells at density of 1×10^6^ cells/ml were stained with 5 µl CM-DiD (Thermo Fisher Scientific) solution and injected into the yolk sac of zebrafish wild-type AB line embryos on day 2 post fertilization. The next day, xenotransplanted embryos bearing labeled MV4-11 cells were treated with 200 nM MA191 or the respective solvent control. Animals were randomly distributed into treatment or control groups. Tumor growth was assessed using an ImageXpress Confocal High-Content Imaging System (Molecular Devices), both before drug exposure (day 1) and 48 hours (day 3) after treatment. The tumor volume was quantified unbiasedly with a semi-automated macro for ImageJ as previously described [33]. Tumor growth was measured by calculating volume differences [%] between day 1 and day 3 imaging points. To determine the responses to the drug, the zebrafish-adapted Response Evaluation Criteria in Solid Tumors (RECIST) was used [33].

### Co-culture of AML cells with bone marrow stromal fibroblasts

HS-5 cells were seeded at a density of 2 x 10^5^ cells/well in a 6-well plate. When cells reached 70% confluency, AML cells were seeded at 1 x 10^5^ per well in 2 ml complete RPMI-1640 medium into thincert cell culture inserts with 0.4 μm pore polyester membrane (657640, Greiner, Frickenhausen, Germany) placed on the HS-5 cells. After 24 h co-culture, cells were treated for indicated time periods, harvested, and analyzed.

### Statistical analyses

Statistics were performed with GraphPad Prism 6. Significant differences between groups were determined by one-way or two-way ANOVA, as indicated. Multiple comparisons were made with Bonferroni correction. Asterisks indicate p values (*p, <0.05; **p, <0.01; ***p, <0.001; ****p, <0.0001). Error bars represent standard error of the mean (SEM).

## Results

### Characterization of the FLT3 PROTAC MA191

Based on our previously described selective FLT3-ITD PROTAC MA49 [29], we developed a series of new FLT3 PROTACs which includes the compound MA191. Like MA49, MA191 comprises the tyrosine kinase inhibitor part MA68 and a VHL-recruiting domain. Promisingly, 50 nM MA191 induces more late apoptosis than 50 nM MA49 in leukemia cells carrying FLT3-ITD after 72 h Abdelsalam, Halilovic, Krämer, and Sippl, manuscript in preparation. The chemical structures of MA49 and MA191 are shown in **Fig. 1a**.

**Figure 1.**
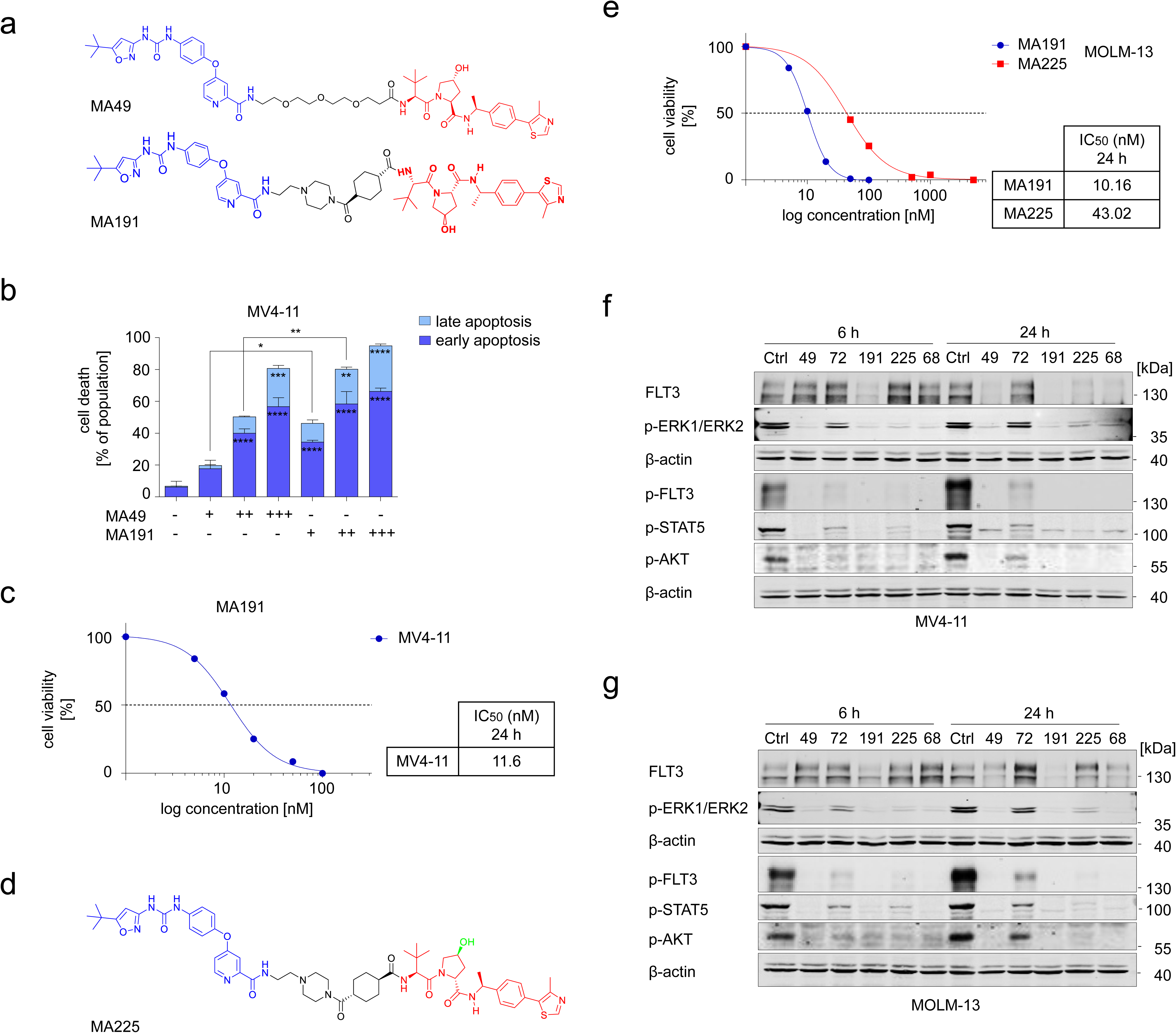
Biological characterization of MA191 FLT3 PROTAC. **a** Structures of MA49 and MA191, blue indicates the FLT3 ligand warhead, red indicates the VHL anchor, black indicates the linker. **b** MV4-11 cells were treated with 10, 20, or 50 nM MA49 or MA191 for 72 h and analyzed for apoptosis by annexin-V and PI staining using flow cytometry. The data are representative for the outcome of three independent experiments; -, untreated, +, 10 nM, ++ 20 nM, +++ 50 nM. Two-way (for early and late apoptosis; asterisks in bars) and one-way (for total apoptosis; asterisks above bars) ANOVA tests were used to determine statistical significance; *p, <0.05; **p, <0.01; ***p, <0.001; ****p, <0.0001. **c** IC_50_ value for apoptosis induction by MA191 in MV4-11 cells was determined by annexin-V and PI staining and flow cytometry. The cells were treated with 0, 5, 10, 20, 50, or 100 nM MA191 for 24 h. IC_50_ values were calculated with GraphPad Prism 6. **d** Structure of MA225, blue indicates FLT3 ligand warhead, red indicates VHL anchor, black indicates the linker, green indicates stereochemical inversion resulting in a VHL inactive stereoisomer. **e** IC_50_ values for apoptosis induction by MA191 and MA225 in MOLM-13 cells were determined by annexin-V and PI staining and flow cytometry. IC_50_ of MA191 was determined as described in **c**. For MA225 IC_50_ determination, MOLM-13 cells were treated with 0.05, 0.1, 0.5, 1, or 5 µM for 24 h. IC_50_ values were calculated with GraphPad Prism 6. **f** MV4-11 and **g** MOLM-13 cells were treated with MA49, MA72, MA191, MA225, or MA68 for 6 h or 24 h. Concentrations used: 50 nM for MA49, MA72, MA191, and MA68; 100 nM for MA225. Lysates of these cells were subjected to immunoblot analyses for FLT3, pT202/pY204-ERK1/ERK2, pY591-FLT3, pY694-STAT5, and pS473-AKT. The protein levels of β-actin were determined to verify equal loading of samples on all tested membranes. The data are representative for the outcome of two independent experiments; phosphorylated; kDa, molecular weight in kilodalton.

We compared how these agents affected leukemic cell survival in more detail. We treated MV4-11 AML cells (carry two copies of FLT3-ITD [34]) with MA49 or MA191 and analyzed apoptosis induction by annexin-V/PI staining followed by flow cytometry. A 72-h treatment with 10-20 nM MA191 caused significantly more apoptosis in MV4-11 cell cultures than the corresponding doses of MA49 (**Fig. 1b**). This notion shows that MA191 is more effective than MA49 at low nanomolar concentrations.

We determined the growth-inhibitory IC_50_ value of MA191 after 24 h and at a concentration range between 0-100 nM as 11.6 nM (**Fig. 1c**). We also calculated the IC_50_ values for an application range of 0-100 nM MA191 for 72 h when applied to MV4-11 and MOLM-13 cells (carry FLT3-ITD and FLT3 [34]). The IC_50_ values were between 9.6 and 11 nM (**Supplementary Fig. S1a**).

We aimed to distinguish the relevance of FLT3-ITD degradation versus FLT3 inhibition by MA191. For this approach, we synthesized a negative control molecule for MA191, named MA225. This agent contains an inactive stereoisomer of the VHL ligand. Consequently, MA225 cannot recruit VHL but has the same inhibitor part as MA191 (**Fig. 1d**). The structures of MA49, MA191, and MA225 are depicted in **Fig. 1a,1d**.

The IC_50_ value of MA225 in MOLM-13 cells was 4-fold higher than the IC_50_ of MA191 (**Fig. 1e**). To study this further, we applied 50 nM MA191 and deliberately applied a higher dose of 100 nM MA225 to MV4-11 and MOLM-13 cells for 6, 16, and 24 h. A concentration of 100 nM of MA225 was necessary to induce apoptosis of these cells comparably as 50 nM MA191 did after 16 h and 24 h (**Supplementary Fig. S1b**). This higher potency of MA191 compared to MA225 verified the advantage of FLT3-ITD degradation over its inhibition.

Next, we analyzed if MA191 and MA225 induce the anticipated breakdown of FLT3-ITD and its downstream signaling in MV4-11 and MOLM-13 cells by immunoblot. As comparators, we used our previously characterized FLT3 PROTAC MA49, its control MA72 that cannot recruit VHL, and MA68, which is the parental FLT3 inhibitor that we used as backbone for the chemical synthesis of our new compounds [29]. To assess molecular processes that precede apoptosis and those that are associated with apoptosis, we treated the cells for 6 h and 24 h. After 6 h treatment, we did not observe apoptosis induction with the very sensitive annexin-V/PI assay (**Supplementary Fig. S1b**). We found that all tested compounds inhibited autophosphorylation of FLT3 (pY591) and phosphorylation of ERK1/ERK2, STAT5 and AKT after 6 h in both MV4-11 (**Fig. 1f**) and MOLM-13 AML cells (**Fig. 1g**). MA49, MA191, and MA68 conferred the highest efficacy in this regard, but only MA191 induced the degradation of FLT3-ITD after 6 h (**Fig. 1f,g**). After 24 h, MA191 had the strongest depleting impact on FLT3-ITD in MV4-11 and MOLM-13 cells. MA225 and MA68 reduced FLT3-ITD in MV4-11 cells but not MOLM-13 cells (**Fig. 1f,g**). This corresponds to a degradation of FLT3-ITD by caspases being the main mediators of apoptosis upon long-term treatment with FLT3 inhibitors [35, 36].

In both AML cell systems, MA49 and MA191 prevented an accumulation of the phosphorylated forms of the MAPKs ERK1 and ERK2 at 24 h. We observed ERK1/ERK2 phosphorylation with the other compounds at 24 h albeit they abrogated this process at 6 h treatment periods (**Fig. 1f,g**). These findings demonstrate that MA49 and MA191 can cease ERK activation, which is known to confer drug resistance [14, 15].

These results identify MA191 as a new nanomolar FLT3 PROTAC that is superior to the previously described MA49. Nanomolar concentrations of MA191 deplete FLT3-ITD rapidly and disengage anti-apoptotic MAPK signaling.

### MA191 is a *bona fide* VHL-based PROTAC that induces BIM-dependent FLT3 degradation

Because the MA49-induced FLT3 degradation was associated with a BIM-dependent depletion of heat shock proteins [29], we investigated if this applied to MA191. We treated MV4-11 cells with 10, 20, or 50 nM MA191 for 16-24 h and analyzed the cells by immunoblot. Within this timeframe, MA191 induced apoptosis up to 40% (**Supplementary Fig. S1b**). MA191 decreased FLT3-ITD and HSP110 dose- and time-dependently (**Fig. 2a**). Concerning HSP110, we noted the occurrence of a cleavage product concomitant with the activation of caspase-3 (**Fig. 2a**). HSP70 is a molecular chaperone necessary for the folding and degradation of VHL E3 ligase [37]. Compared to FLT3-ITD and HSP110, HSP70, HSP27, and HSP90 were reduced to a lesser extent by MA191. In addition, MA191 entirely inhibited the phosphorylation of FLT3-ITD at all analyzed treatment periods and attenuated phosphorylated STAT5 dose- and time-dependently. A time- and dose-dependent accumulation of active caspase-3 subunits started to occur after 16 h with 20 nM MA191. Concomitant herewith, BIM-EL, the major splice variant of pro-apoptotic BIM, began to accumulate after a 16 h incubation with 20 nM and 50 nM MA191 (**Fig. 2a**).

**Figure 2.**
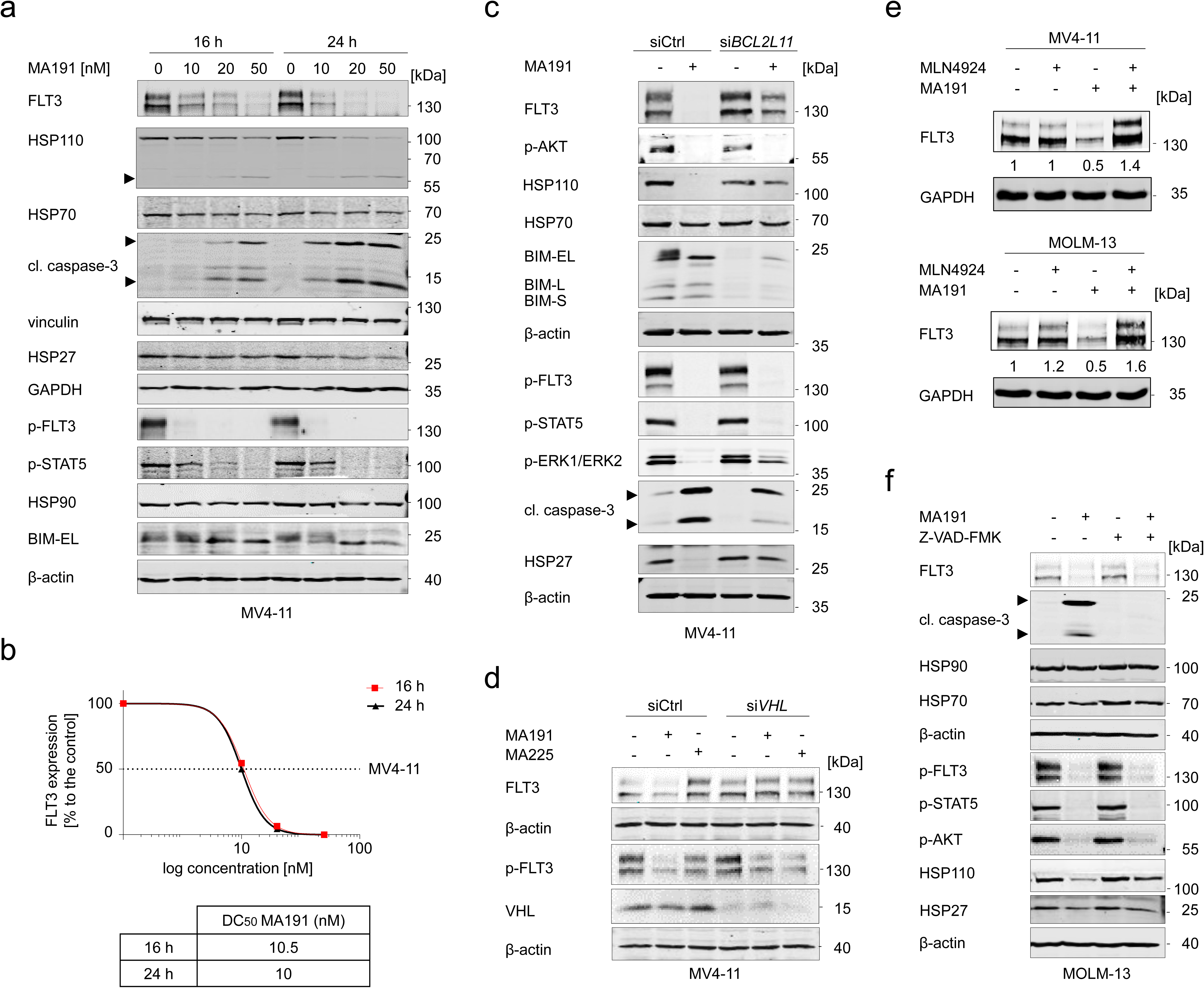
MA191 is BIM-dependent VHL PROTAC. **a** MV4-11 cells were treated with 10, 20, or 50 nM MA191 for 16 h or 24 h and analyzed by immunoblot for the expression of FLT3, HSP110, HSP90, HSP70, HSP27, cleaved caspase-3, pY591-FLT3, pY694-STAT5, and BIM-EL. The protein levels of vinculin and β-actin were determined to verify the equal loading of samples. The data are representative for the outcome of three independent experiments; cl., cleaved form, arrows point to the cleavage fragments of active caspase-3; p-, phosphorylated; EL, extra-long splice form; kDa, molecular weight in kilodalton. **b** DC_50_ values for FLT3 degradation by MA191 in MV4-11 cells were determined by quantitative immunoblot using the Odyssey system. The cells were treated as stated in **a** and lysates were analyzed for FLT3 and vinculin. The values for FLT3 were normalized to vinculin and the DC_50_ values were calculated with GraphPad Prism 6. **c** MV4-11 cells were transfected with siRNA against the *BCL2L11* mRNA to knock down the BIM protein. Cells with control siRNA and with the BIM knockdown were treated with 50 nM MA191 for 24 h and analyzed by immunoblot for FLT3, pT202/pY204-ERK1/ERK2, pY591-FLT3, pY694-STAT5, pS473-AKT, the three BIM splice variants extra-long (EL), long (L), short (S), cleaved caspase-3, HSP110, HSP90, HSP70, and HSP27. The protein levels of β-actin were determined to verify equal loading of samples. The data are representative for the outcome of two independent experiments; +, treated; -, untreated; p-, phosphorylated; kDa, molecular weight in kilodalton; siCtrl, siRNA control; si*BCL2L11*, siRNA targeting BIM; cl., cleaved, arrows point to the cleavage fragments of active caspase-3. **d** MV4-11 cells with control siRNA or with a VHL knockdown were treated with 50 nM MA191 or 100 nM MA225 for 6 h. Lysates of the cells were analyzed by immunoblot for FLT3, pY591-FLT3, and VHL. The protein levels of β-actin were determined to verify equal sample loading. The data are representative of the outcome of two independent experiments; +, treated; -, untreated; p-, phosphorylated; kDa, molecular weight in kilodalton; siCtrl, siRNA control; si*VHL*, VHL knockdown by RNAi. **e** MV4-11 and MOLM-13 cells were treated with 50 nM MA191 ± 3 µM MLN4924 for 6 h and analyzed by immunoblot for FLT3. The protein levels of GAPDH were determined to verify equal loading of samples. The data are representative for the outcome of two independent experiments in each cell line; +, treated; -, untreated; kDa, molecular weight in kilodalton. **f** MOLM-13 cells received a pre-treatment with Z-VAD-FMK for 1 h if indicated (+). The cells were treated with 50 nM MA191 for 24 h and analyzed by immunoblot for FLT3, cleaved caspase-3, pY591-FLT3, pY694-STAT5, pS473-AKT, HSP110, HSP90, HSP70, and HSP27. The protein levels of β-actin were determined to verify equal loading of samples. The data are representative of the outcome of two independent experiments; +, treated; -, untreated; p-, phosphorylated; kDa, molecular weight in kilodalton; cl., cleaved, arrows point to the activated cleavage fragments of caspase-3.

Since BIM is a key regulator of intrinsic apoptosis that occurs upon lost mitochondrial integrity [29], we measured mitochondrial membrane potentials of untreated and MA191-treated MV4-11 cells. After a 24 h application, MA191 evoked a breakdown of the mitochondrial membrane potential (ΔΨm). This was not evident after a 6 h treatment (**Supplementary Fig. S1c**). These data support that ΔΨm is associated with apoptosis and they corroborate that the rapid depletion of FLT3-ITD by MA191 is not due to apoptosis-associated events (**Fig. 1f,g, Supplementary Fig. S1c**).

As expected, FLT3-ITD was more affected by MA191 than the other proteins that we analyzed (**Fig. 2a**). Since MV4-11 cells are homozygous for FLT3-ITD, we could use this cell line to calculate the half-maximal degradation concentrations (DC_50_) of FLT3-ITD in the presence of MA191. The DC_50_ values of MA191 are 10.5 nM for 16 h and 10 nM for 24 h treatment periods (**Fig. 2b**).

To study the role of BIM in mediating the effects of MA191, we transfected MV4-11 cells with siRNA targeting the BIM-encoding mRNA *BCL2L11*. We confirmed the knockdown (KD) efficiency for BIM by immunoblot as reflected by downregulation of its three splice variants BIM-EL, BIM-L, and BIM-S. The KD of BIM suppressed the MA191-induced degradation of FLT3, which was linked to stable HSP110 and HSP27 levels in MA191-treated MV4-11 cells. The MA191-mediated inhibition of the phosphorylation of FLT3-ITD, AKT, and STAT5 remained unaffected by KD of BIM (**Fig. 2c**). This finding shows that BIM controls the reduction of FLT3-ITD by MA191 but not the inhibition of its catalytic activity by the FLT3 inhibitor moiety of MA191.

The suppression of FLT3-ITD and its downstream signaling cascade by MA191 in MV4-11 cells with a KD of BIM can explain the not fully repressed activation of caspase-3. Inhibited FLT3-ITD can activate MAPK signaling from the cell surface [14, 15]. Thus, the inhibition of ERK1/ERK2 phosphorylation by MA191 should be partially rescued in cells with BIM KD and stabilized FLT3-ITD. Immunoblot analyses verified this hypothesis (**Fig. 2c**).

To confirm the VHL-based mode of activity of MA191, we applied siRNA targeting the *VHL* mRNA to MV4-11 cells and subsequently incubated them with MA191 or MA225 for 6 h. Immunoblot verified the VHL KD. We noted that the MA191-induced degradation of FLT3 was rescued in cells with a KD of VHL (**Fig. 2d**). This demonstrates that VHL is necessary for the MA191-mediated degradation of FLT3. The reduction of FLT3 autophosphorylation after MA191 treatment in cells with a VHL KD was comparable to the impact of FLT3 inhibition by MA225 (**Fig. 2d**). This observation illustrates that MA191 acts as a FLT3 inhibitor in the absence of VHL but lacks degrading properties on FLT3-ITD.

Having shown that MA191 requires BIM and VHL to deplete FLT3-ITD, we aimed to confirm the anticipated role of the CRL2^VHL^ E3 ubiquitin ligase complex [16, 38]. Neddylation is a process in which the ubiquitin-like protein NEDD8 is covalently attached to the target protein. This posttranslational modification drives CRL2^VHL^ to undergo conformational changes and multivalent interactions that accelerate the ubiquitination of substrate proteins [38]. Thus, the role of CRL2^VHL^ can be studied using the neddylation inhibitor MLN4924. We applied MLN4924 in combination with MA191 to MV4-11 and MOLM-13 cells. MLN4924 prevented the MA191-mediated degradation of FLT3-ITD in both AML cell types (**Fig. 2e**).

To exclude the possibility that MA191 depletes FLT3-ITD due to apoptosis-activated proteolysis, we treated MOLM-13 cells with MA191 and the pan-caspase inhibitor Z-VAD-FMK for 24 h and analyzed cell lysates by immunoblot. The MA191-induced caspase-3 cleavage was counteracted entirely by Z-VAD-FMK, confirming full apoptosis inhibition. Irrespective thereof, MA191 remained a potent PROTAC against FLT3-ITD in MOLM-13 cells that were incubated with Z-VAD-FMK. MA191 degraded FLT3 and inhibited its phosphorylation, as well as the phosphorylation of its downstream targets STAT5 and AKT (**Fig. 2f**). Hence, MA191 depletes FLT3-ITD independently of caspase activity.

We additionally analyzed HSPs to examine if their downregulation by MA191 was associated with apoptosis activation. HSP70, HSP110, and HSP27 were stabilized in cells pre-treated with Z-VAD-FMK, suggesting that their downregulation is mediated by caspases (**Fig. 2f**).

These results confirm that MA191 is a potent VHL-based PROTAC that requires BIM, and neddylation-dependent proteasomal degradation to eliminate FLT3-ITD.

### MA191 induces apoptosis of cultured and primary AML cells with FLT3-ITD

Using the data resource Hemap (http://hemap.uta.f/), we assessed *FLT3* mRNA expression in various leukemia subtypes and normal hematopoietic cells. Compared to normal erythroid, myeloid, and lymphoid lineages, *FLT3* was expressed at higher levels in acute lymphoblastic leukemia (ALL) and AML cells (**Fig. 3a**). According to the DepMap Portal (https://depmap.org/), B-lymphoblastic leukemia/lymphoma and AML cell lines, particularly MV4-11 and MOLM-13, are highly dependent on FLT3 (**Fig. 3b**). This makes MV4-11 and MOLM-13 cells most suitable for studying FLT3-targeting drugs.

**Figure 3.**
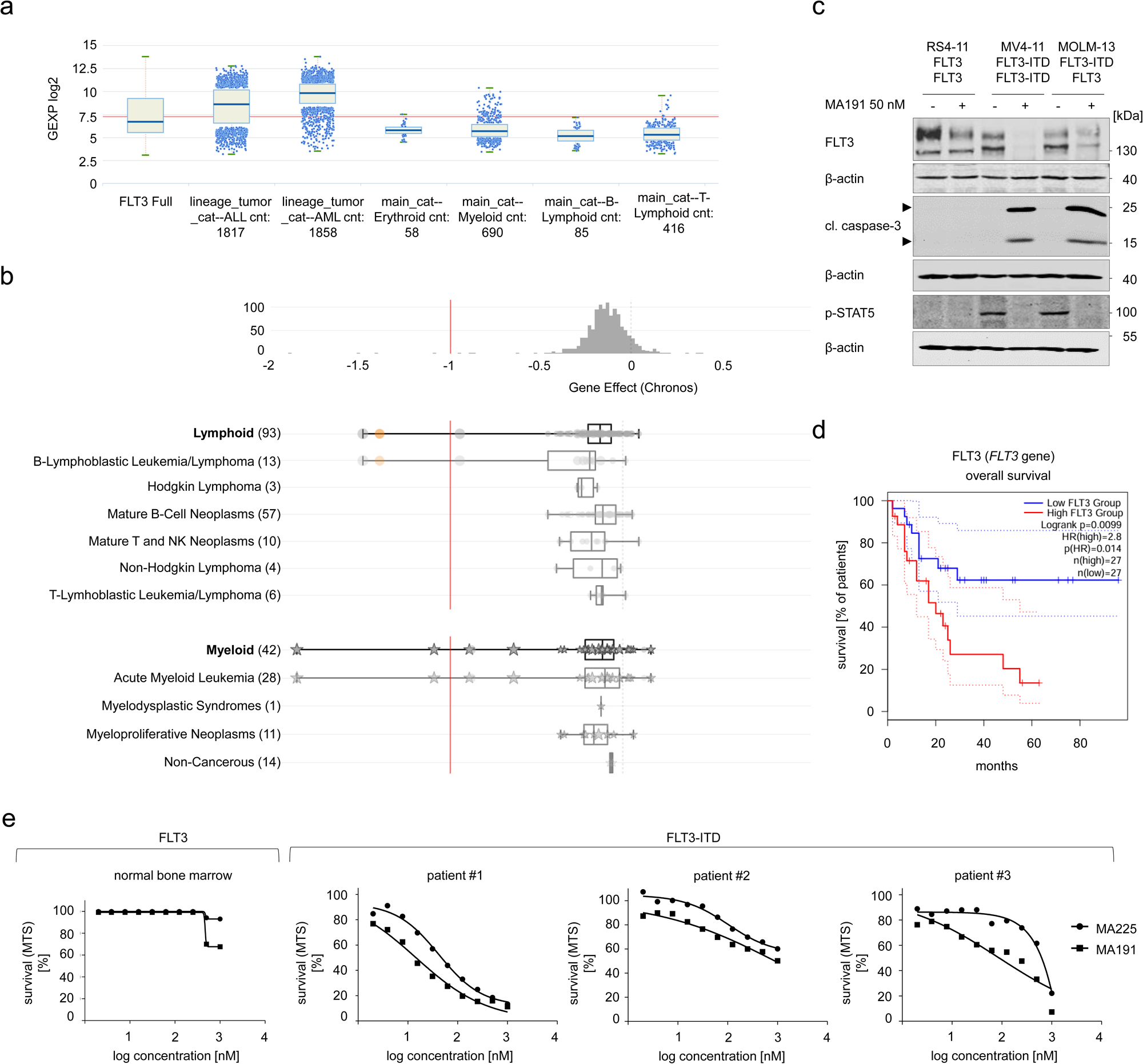
MA191 targets cultured and primary AML cells with mutant FLT3. **a** Gene expression of *FLT3* mRNA in acute lymphoblastic leukemia (ALL), acute myeloid leukemia (AML), and normal erythroid, myeloid, B-lymphoid, and T-lymphoid cells. Data was retrieved from HEMAP: Online resource for interactive exploration and e-staining of hematopoietic cancer data. **b** Analysis of the dependency of lymphoid and myeloid cancer cells on FLT3 according to data obtained from the DepMap database. **c** RS4-11, MV4-11, and MOLM-13 cells were treated with 50 nM MA191 for 24 h. Cell lysates were subjected to immunoblot analyses for FLT3, cleaved caspase-3, pY694-STAT5, and pS473-AKT. The protein levels of β-actin were determined to verify equal loading of samples. The data are representative for the outcome of three independent experiments; p-, phosphorylated; kDa, molecular weight in kilodalton; cl., cleaved, arrows point to the cleavage fragments of active caspase-3. **d** The GEPIA2 database was analyzed for a correlation between overall survival of AML patients and the levels of FLT3. The analysis included 54 patients in total, with 50% in the high and 50% in the low expressing groups (p=0,014). **d** Serial dilutions of 50 µM (#3; top concentration; decreasing in log) of MA191 or MA225 were applied to human primary normal bone marrow and primary samples from leukemia patients with FLT3-ITD for 72 h. AML patient samples are designated as #1, #2, and #3. Cell survival is provided as %-values compared to untreated control cells (100% survival) in the MTS-assay.

Considering these findings, we assessed the impact of MA191 on FLT3 versus FLT3-ITD. We compared MV4-11, MOLM-13, and RS4-11 (ALL) cells, which express wild-type FLT3. We treated them with MA191, analyzed apoptosis induction by flow cytometry, and probed immunoblots with an antibody against FLT3. In MV4-11 and MOLM-13 cells, MA191 decreased the FLT3-ITD levels after 24 h, whereas in RS4-11, FLT3 was less affected. Furthermore, we noted a FLT3-variant-specific decline, with heterozygous FLT3-ITD being the most and FLT3 being less degraded by MA191 (**Fig. 3c**). These results suggest that MA191 has a higher affinity for FLT3-ITD compared to FLT3. Since FLT3-ITD-expressing AML cells critically depend on this oncogene, we observed an induction of apoptotic cleavage of caspase-3, as well as the inhibition of the activating downstream phosphorylation of STAT5 specifically in MV4-11 and MOLM-13 cells. RS4-11 cells showed no detectable activation of STAT5 (**Fig. 3c**). Flow cytometry consistently demonstrated that MA191 did not induce apoptosis in RS4-11 after a 72-h treatment. In MV4-11 cell cultures, the death rate was nearly 100% and in the case of MOLM-13 cell cultures it was 50% (**Supplementary Fig. S2a**).

To assess the specificity of MA191 further, we evaluated its impact on c-KIT. This class III RTK structurally resembles FLT3 [6]. We used HMC-1.1 and HMC-1.2 cells that have a V560G KIT mutation or a double V560G and D816V KIT mutations. Growth of these cell lines requires constitutively active c-KIT [39]. The application of 100 nM MA191 induced 22% apoptosis in HMC-1.1 cells after 72 h. HMC-1.2 cells were insensitive to MA191 at all tested concentrations (**Supplementary Fig. S2b**). Even though there was an apoptosis induction in HMC-1.1 cells, 50-100 nM MA191 did not reduce c-KIT levels. However, our finding that MA191 decreased phosphorylated c-KIT in HMC-1.1 cells (**Supplementary Fig. S2c**) may explain the observed growth inhibitory effects (**Supplementary Fig. S2b**).

Survival analysis data that we retrieved from the Gene Expression Profiling Interactive Analysis (GEPIA2) database (http://gepia2.cancer-pku.cn/) showed significantly lower survival for AML patients with high compared to low FLT3 expression (**Fig. 3d**). These data may hint to a benefit of drugs that attenuate the levels of FLT3.

We next tested how the depletion of FLT3-ITD by MA191 versus the inhibition of FLT3-ITD by MA225 affected primary AML patient samples with FLT3-ITD. As a comparator, we tested normal human bone marrow samples. Both MA191 and MA225 reduced the survival of FLT3-ITD-positive AML cell samples, with MA191 having lower EC_50_ values than MA225 (**Fig. 3e**). Information on patients regarding their FLT3 and NPM1 mutation status, and EC_50_ values of MA191 and MA225 is denoted in **Table 2**. NPM1 mutational status had no impact on AML cell responses to MA191 and MA225. Normal bone marrow cells were only affected at 50 µM doses of MA191 and MA225 (**Fig. 3e**).

These data illustrate that FLT3 is dysregulated in leukemia cells, associated with worse prognosis, and shows high cytotoxic selectivity for AML cells with FLT3-ITD.

### MA191 does not affect normal immune cells, hematopoietic stem cells, and progenitor cells

The Human Protein Atlas database (https://proteinatlas.org/) reports for hematopoietic cell types that the *FLT3* mRNA is in most highly expressed in myeloid and plasmacytoid dendritic cells, followed by hematopoietic progenitor cells and monocytes (**Fig. 4a**).

**Figure 4.**
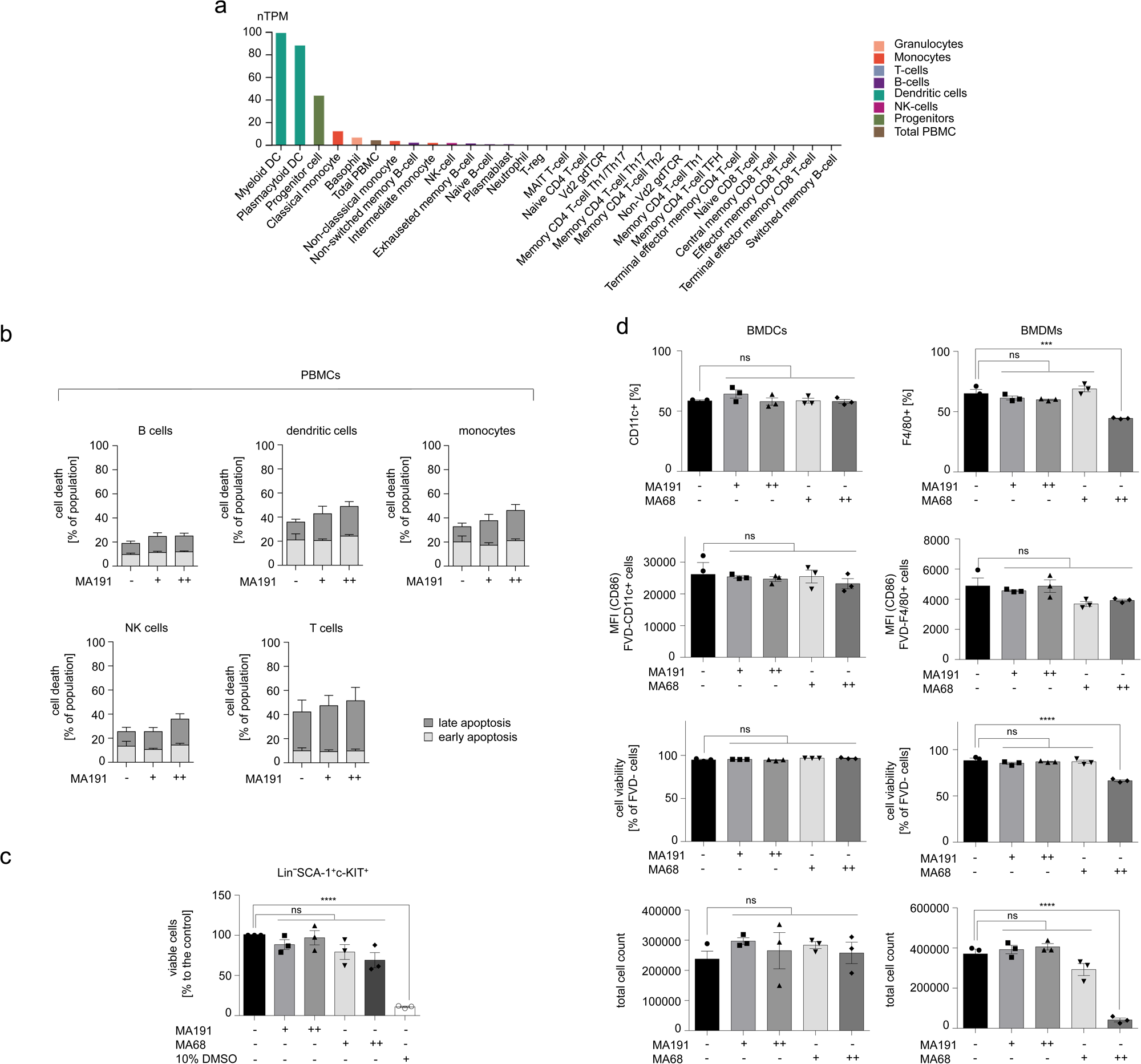
MA191 is not toxic to normal immune and hematopoietic stem cells and has in vivo activity. **a** *FLT3* transcript expression values in nTPM, according to the internal normalization pipeline for 18 immune cell types and total peripheral mononuclear cells (PBMC); data was retrieved from The Human Protein Atlas database. **b** Apoptosis analysis of PBMC subpopulations after treatment with 20 or 50 nM of MA191 for 48 h; PBMC, peripheral blood mononuclear cells; NK cells, natural killer cells; -, untreated, +, 20 nM, ++ 50 nM. The data are representative for the outcome of four independent experiments. Two-way ANOVA statistical test was used to determine statistical significance. **c** Cell viability analysis of murine bone marrow stem cells. Bone marrow tissues were collected from the hind legs of previously killed C57BL/6 mice. The cells were treated with 50 or 100 nM of MA191, MA68, or a high concentration of 10% DMSO as a positive control for cytotoxicity for 24 h; -, untreated, +, 50 nM, ++ 100 nM. One-way ANOVA statistical test was used to determine statistical significance; ns, not significant; *p, <0.05; **p, <0.01; ***p, <0.001; ****p, <0.0001; n=3, male and female. **d** Bone marrow cells were isolated from C57BL/6 mice and cultured for 7 days in medium containing the inhibitors at 50 or 100 nM concentration and supplementation with either GM-CSF to induce the differentiation of myeloid progenitor cells to BMDCs or M-CSF to induce the differentiation to BMDMs. After one week, we assessed the total cell number, viability and frequencies of CD11c+ (pan dendritic cell marker) or F4/80+ (macrophage marker) cells, the expression of their activation marker CD86, cell viability (negative for FVD), and the total cell numbers; -, untreated, +, 50 nM, ++ 100 nM. One-way ANOVA statistical test was used to determine statistical significance; ns, not significant; *p, <0.05; **p, <0.01; ***p, <0.001; ****p, <0.0001; n=3, male and female.

Next, we asked if MA191 impairs the survival of human peripheral blood mononuclear cells (PBMCs). We discriminated B cells, dendritic cells, monocytes, NK cells, and T cells by flow cytometry assessing cell type-specific surface markers. We treated these cell populations with 20 or 50 nM MA191 for 48 h and analyzed them for apoptosis induction. Neither of the tested concentrations affected the viability of normal immune cells significantly (**Fig. 4b**).

Next, we assessed the effect of MA191 on hematopoietic stem cells that were isolated from murine bone marrow. We treated these cells with 50 or 100 nM MA191 or MA68 for 24 h. DMSO at 10% concentration served as positive control for toxicity. A treatment with 50-100 nM MA191 did not compromise the viability of murine stem cells. MA68 had a detectable but non-significant effect on the stem cells (**Fig. 4c**).

We also evaluated how 50-100 nM MA191 affected the differentiation of hematopoietic stem cells of the myeloid lineage. Isolated murine hematopoietic progenitor cells were supplemented with GM-CSF or macrophage colony-stimulating factor (M-CSF) to yield bone marrow-derived dendritic cells (BMDCs) or bone marrow-derived macrophages (BMDMs), respectively. We treated the cells with either MA191 or MA68 for seven days and then evaluated for total cell count, cell viability, frequencies of CD11c^+^ (dendritic cell marker) or F4/80^+^ (macrophage marker) cells, and the expression of their common activation marker CD86. Both concentrations of MA191 did not affect the survival, proliferation, and differentiation of BMDCs and BMDMs (**Fig. 4d**). This finding verifies its safety for hematopoietic cell differentiation. Although MA68 did not compromise the differentiation of BMDCs, it hampered the proliferation, cell viability, and differentiation of BMDMs significantly (**Fig. 4d**).

We conclude that MA191 does not disrupt normal immune cells, hematopoietic stem cells, and blood cell maturation, and that MA191 is superior to its cognate kinase inhibitor.

### MA191 is effective in vivo and breaks mechanisms of FLT3 inhibitor resistance of AML cells

We consequently studied if MA191 is active against AML cells in vivo. We injected MV4-11 cells into *Danio rerio* (zebrafish) early larvae, let tumor masses establish, and subsequently treated them with MA191 for 48 h. We evaluated the effects of MA191 based on the comparison of leukemia cell masses before and after treatment. Using zebrafish-adapted response evaluation criteria in solid tumors (RECIST), zebrafish larvae can be classified to have developed progressive disease (PD), stable disease (SD), or a partial response (PR) to the treatment. In the solvent-treated group, 72.7% of larvae developed PD and 27.3% developed SD, with 0% of larvae responding to the treatment. In the MA191 group, the rate of larvae with PD decreased to 44%, SD increased to 44%, and 12% of larvae experienced PR (**Fig. 5a**).

**Figure 5.**
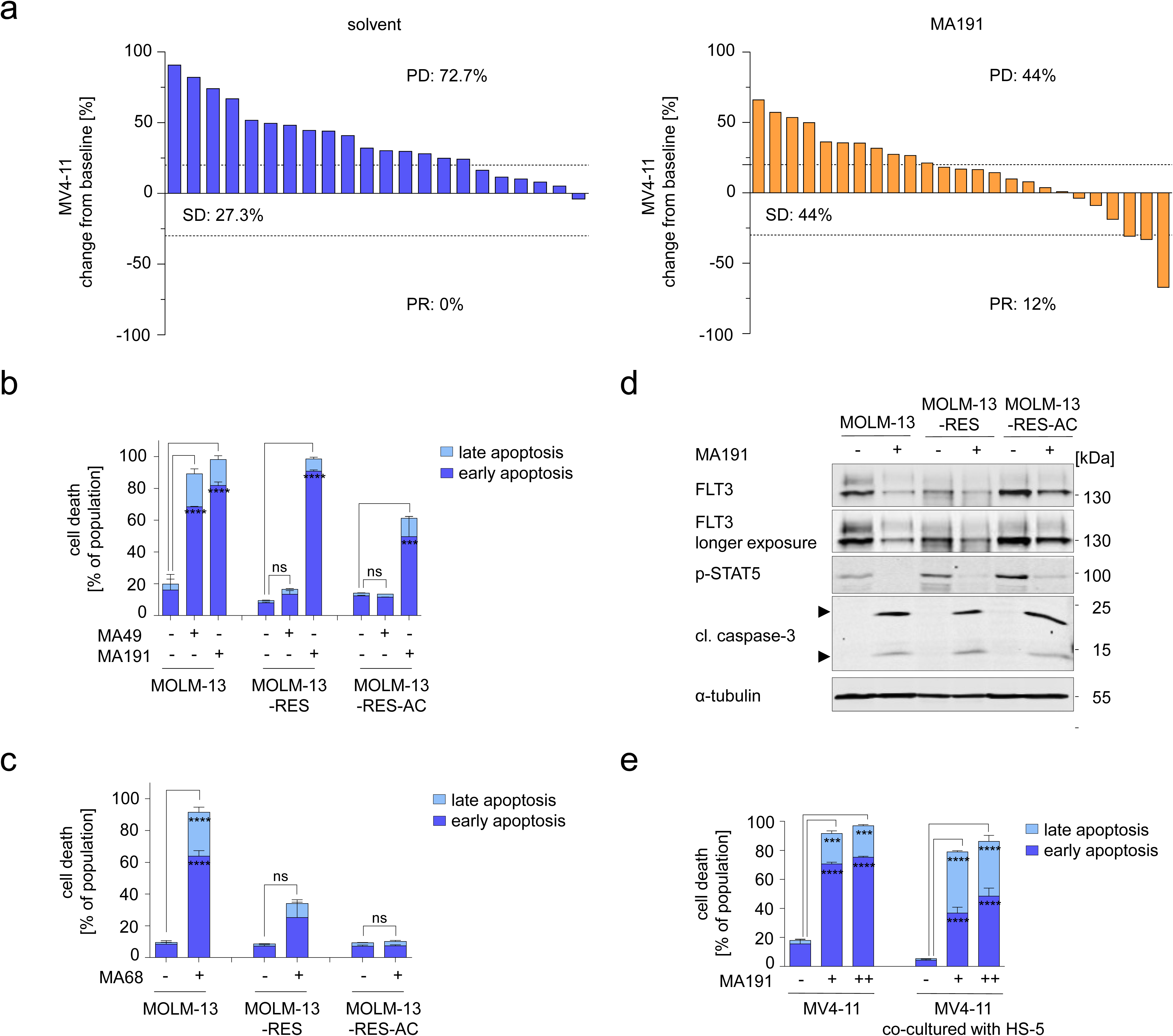
MA191 overcomes extrinsic and intrinsic resistance mechanisms of AML. **a** Waterfall plots show changes in tumor volume [%] for each individual zebrafish larvae engrafted with MV4-11 cells, from baseline (day 1 = start of the treatment) until day 3 after cell injections. Zebrafish larvae xenografts were treated with DMSO used as a solvent (n=22 larvae; left) or 200 nM MA191 (n=25 larvae; right) for 48 h; each bar reflects one individual xenograft. Numbers indicate the percentage of early larvae with progressive disease (PD), stable disease (SD) and partial response (PR) in each treatment group on day three. **b,c** MOLM-13 (with one FLT3-ITD allele), MOLM-13-RES (with secondary D835Y mutation on FLT3-ITD allele), and MOLM-13-RES-AC (homozygous for secondary D835Y mutation on two FLT3-ITD alleles), were treated with 200 nM MA49, MA191 (**b**), or 200 nM MA68 (**c**) for 72 h and analyzed for apoptosis by annexin-V and PI staining using flow cytometry. The data are representative for the outcome of two independent experiments; -, untreated, +, 200 nM. Two-way ANOVA statistical test was used to determine statistical significance; ns, not significant; *p, <0.05; **p, <0.01; ***p, <0.001; ****p, <0.0001. **d** MOLM-13, MOLM-13-RES, and MOLM-13-RES-AC were treated with 200 nM MA191 for 24 h. Cell lysates were analyzed by immunoblot for the expression of FLT3, cleaved caspase-3, pY694-STAT5, and BIM. The protein levels of α-tubulin were determined to verify equal loading of samples. The data are representative for the outcome of three independent experiments; -, untreated, +, 200 nM; p-, phosphorylated; kDa, molecular weight in kilodalton; cl., cleaved, arrows point to the cleavage fragments of active caspase-3. **e** MV4-11 cells without and with co-cultured bone marrow stromal fibroblast cell line HS-5 were treated with 50 or 100 nM MA191 for 72 h and analyzed for the apoptosis by annexin-V and PI staining using flow cytometry. MV4-11 cells were co-cultured with HS-5 cells for 24 h prior the treatment. The data are representative for the outcome of three independent experiments; -, untreated, +, 50 nM, ++, 100 nM. Two-way ANOVA statistical test was used to determine statistical significance; *p, <0.05; **p, <0.01; ***p, <0.001; ****p, <0.0001

Secondary mutations in the FLT3 TKD undermine the therapeutic management of AML with FLT3-ITD. This raises a need for effective therapeutics that break such resistance mechanisms. We speculated that the ability of MA191 to deplete FLT3-ITD could be a strategy to eliminate FLT3-ITD/TKD mutants. We applied MA191 to cells with acquired D835Y mutations on FLT3 alleles with ITD mutations. We used the MOLM-13-RES cell line with an acquired D835Y TKD mutation on one of its two FLT3-ITD alleles or the MOLM-13-RES-AC cell line with D835Y TKD mutations on both FLT3-ITD alleles (the cells are described in [30]). We treated MOLM-13, MOLM-13-RES, or MOLM-13-RES-AC with MA191 or MA49 and analyzed apoptosis induction after 48 h (**Supplementary Fig. S3a**) or 72 h (**Fig. 5b**). Even though MA49 was highly effective against MOLM-13 cells, it did not cause apoptosis in MOLM-13-RES and in MOLM-13-RES-AC cells. MA191 induced significant apoptosis in all tested cell lines and at both assessed time points (**Supplementary Fig. S3a, Fig. 5b**). To complement these analyses, we scrutinized if MA68 targets cells with FLT3-ITD having secondary TKD mutations. MA68 induced cell death in MOLM-13-RES after 48 h (**Supplementary Fig. S3b**) and 72 h (**Fig. 5c**), but the effect was not significant. MOLM-13-RES-AC cell line did not respond to the MA68 treatment (**Supplementary Fig. S3b, Fig. 5b**).

To confirm the outcome of these experiments, we evaluated if MA191 evoked the anticipated elimination of FLT3-ITD/TKD molecules. MA191 reduced FLT3-ITD and FLT3-ITD/TKD in the tested cell lines after 24 h. The degradation of FLT3-ITD/TKD in MOLM-13-RES-AC cells appeared less pronounced. This may stem from higher basal levels of FLT3-ITD/TKD (**Fig. 5d**). Such remaining FLT3-ITD/TKD can explain lower apoptosis induction in MOLM-13-RES-AC cells than in MOLM-13 and MOLM-13-RES cells (**Fig. 5b**). The attenuation of FLT3-ITD and FLT3-ITD/TKD translated into an inhibition of STAT5 phosphorylation. Apoptosis-associated caspase-3 cleavage appeared after 24 h in all tested cell lines (**Fig. 5d**).

Another resistance mechanism of FLT3-ITD-driven AML relies on a protective microenvironment of BM cells [7–10]. To assess whether this protective mechanism disables MA191, we used human bone marrow stromal cell line HS-5 and co-cultured these cells with MV4-11 AML cells before and during the treatment. These cocultures received 50 to 100 nM MA191 for 72 h. We used MV4-11 cells cultured alone in parallel as reference. MA191 induced significant apoptosis in both co-cultured and non-co-cultured MV4-11 cells at all tested concentrations (**Fig. 5e**). However, it did not affect the viability of HS-5 cells (**Supplementary Fig. S3c**), which is consistent with the results obtained for primary normal human cells (**Fig. 4**).

These results show that MA191 is safe and effective in an in vivo *Danio rerio* larvae model and that MA191 is more potent than MA49 and their parent inhibitor MA68. Moreover, MA191 can overcome resistance mechanisms that stem from secondary FLT3 TKD mutations and from the BM microenvironment.

## Discussion

In this study, we characterize anti-leukemia effects of the novel FLT3 PROTAC MA191. This VHL- and MA68-derived PROTAC is more effective than our previously identified VHL- and MA68-derived FLT3 degrader MA49 against human FLT3-ITD-dependent AML cells. MA191 eliminates FLT3-ITD more rapidly and effectively than MA49. This results in a more potent apoptosis induction by MA191.

Various drugs induce the elimination of FLT3-ITD by proteasomes, autophagy, and caspase-mediated cleavage [35, 40, 41]. This work provides evidence that MA191 induces a VHL-dependent depletion of FLT3-ITD that is not just a cause of apoptosis. The detectable elimination of FLT3-ITD by MA191 occurs within few hours and precedes a measurable cell death induction by this agent. The depletion of VHL by RNAi antagonizes the MA191-induced loss of FLT3-ITD. In addition, our results show that the depletion of FLT3-ITD by MA191 relies on neddylation. Since cullin proteins require neddylation for their activation, our findings suggests that MA191 targets FLT3 by recruiting the CRL2^VHL^ complex. As VHL itself is a neddyation substrate, addition of NEDD8 to VHL could be required for the biological effects of MA191 [38, 42].

The FLT3 ligand warhead and the VHL anchor domains of MA49 and MA191 are identical. Therefore, the replacement of the PEG linker of MA49 with a more rigid piperazine linker in MA191 appears as the difference that reasons the higher potency of MA191. The choice of PROTAC linkers can likewise affect PROTAC solubility, cell permeability, efflux, and metabolism. More rigid linkers in PROTACs, containing nitrogen heterocycles instead of traditional PEG and alkyl linkers, also yielded more stable ternary complexes and more effective target elimination in other systems [43, 44]. Thus, more rigid linker structures could become useful in further studies that aim to develop PROTACs.

MA191 and related molecules seem useful to eradicate FLT3-dependent AML cells that are resistant to clinically used FLT3 inhibitors. Furthermore, unlike MA68 and MA49, MA191 can induce apoptosis of AML cells with FLT3-ITD/FLT3-TKD double mutants. This is relevant because clinical success of FLT3 inhibitors is frequently limited by the acquisition of FLT3-TKD mutations in FLT3-ITD [45]. TKD mutations in FLT3 structurally disfavor the binding of type II FLT3 inhibitors that are active in low nanomolar concentrations. Type I inhibitors are less specific and require higher concentrations, that are associated with more undesired side-effects, to achieve anti-leukemic effects [46]. MA68 is based on sorafenib to which the MV4-11 cells that we used are resistant. Thus, it is also a type II inhibitor and is unable to bind FLT3 with TKD mutations. Mild apoptosis induction of MOLM-13-RES cells by MA68 may be due to the presence of one wild-type FLT3 allele in these cells to which MA68 binds. However, MA68 was completely ineffective in MOLM-13-RES-AC cells bearing only FLT3-ITD with the FLT3-TKD D835Y mutation [30]. Unlike conventional small-molecule inhibitors that need to bind to its target’s active site to inhibit enzymatic functions [20–22], PROTACs can act against their targets when they get into close proximity. Thus, a molecular explanation for the efficacy of MA191 against FLT3-ITD/FLT3-TKD is the recruitment of VHL and the associated ubiquitination machinery to these mutants even when the FLT3 ligand warhead of a PROTAC (in our case MA68) is ineffective. The different linkers of MA49 and MA191 provide again an explanation for the formation of a stable ternary complex that supports an effective ubiquitin-mediated proteasomal degradation and resulting superiority of MA191 against AML cells harboring FLT3-ITD/FLT3-TKD. We anticipate that MA191 will also be effective against recently identified transforming FLT3 mutants with deletions in the FLT3 juxtamembrane domain. Patients with such AML cells have a similar sequela as FLT3-ITD-positive patients [47].

MA191 effectively induced apoptosis of AML cells that were co-cultured with human bone marrow stromal fibroblasts. These support the proliferation of hematopoietic progenitor cells via secretion of over 50 pro-survival cytokines, including G-CSF, GM-CSF, M-CSF, IL-1α, IL-1, IL-1RA; IL-6; IL-8; IL-11, and FGF2 [48]. FGF2 overexpression in the bone marrow is one of the most common resistance mechanisms in AML [49]. Our finding that primary and cultured bone marrow cells are insensitive to MA191 suggests its low toxicity. The mitogen-activated kinase-14, also called p38α, is a mediator of the FLT3 inhibitor resistance of FLT3-ITD-expressing AML stem cells that grow on feeder layer cells [50]. It is possible that the co-targeting of FLT3-ITD and p38α by MA191 (Sippl and Krämer, manuscript in preparation or the preprint) disables anti-apoptotic effects of microenvironmental protection signals.

Database analyses show that leukemia cells have higher levels of FLT3 than normal cells and that high FLT3 levels are associated with reduced overall survival. Our finding that MA191 attenuates wild-type FLT3 may point to an option to disable resistance mechanisms that rely on an increased expression of wild-type FLT3 [12]. Although AML patients with FLT3-ITD benefit from quizartinib more than patients with wild-type FLT3, it yielded some beneficial responses in such patients, as well [51]. Therefore, a reduction of FLT3 may be beneficial in a clinical setting.

Despite the strong impact of MA191 on AML cells with FLT3-ITD, MA191 does not affect human PBMCs, murine bone marrow progenitor cells, and leukemia cells with wild-type FLT3. Curiously, MA191 affects the activation of wild-type FLT3 without compromising cell growth. This corresponds to the FLT3-independent growth of such cells. The ability of MA191 to target FLT3 may though overcome a therapy-induced upregulation of the FLT3 ligand that activates compensatory FLT3 signaling in AML cells [4, 49].

The comparative analysis of the PROTAC MA191 and its cognate inhibitor MA225 demonstrates that the elimination of FLT3-ITD is more effective than its sole inhibition. Because of their high similarity, these stereoisomers are optimal molecules for such analyses. The stereoisomers share the same atoms, import, export, and elimination mechanisms. A logical explanation for the improved potency of MA191 is that the removal of FLT3-ITD is more pronounced than its reversible inhibition. The proteasomal degradation of FLT3-ITD necessitates the AML cells to transcribe the *FLT3-ITD*-coding mRNA and to synthesize FLT3-ITD de novo. This mechanism of action may allow a less frequent application of targeted protein degraders in a therapeutic setting. Proton pump inhibitors for gastric disorders have realized such a principle of prolonged inhibition by inactivation for decades [53].

The divergent effects of FLT3 inhibitors and PROTACs on signaling molecules and protein stability can also explain the more potent activities of FLT3 PROTACs. The activation of ERK1/ERK2 by inhibited FLT3-ITD molecules that translocate to the plasma membrane can blunt the apoptotic effects of FLT3 inhibitors [14]. Such an activation of ERK is disabled when PROTACs eliminate FLT3-ITD. It is possible that the superior effects of FLT3 PROTACs result from their impact on HSPs (this work and [29]). MA49 and MA191 pronouncedly reduce HSP110 which allows the proper folding and recycling of proteins [54]. It will be interesting to see if the loss of such a major HSP causes an accumulation of cytotoxic protein aggregates. Because the pan-caspase inhibition rescued HSP110 from MA191-mediated downregulation and due to the advent of a HSP110 cleavage product, we deduce that the loss of HSP110 is an apoptosis-associated event. The EXPASY peptide cutter indicates a caspase-1 cleavage site in HSP110 at amino acid position 150 (https://web.expasy.org/cgi-bin/peptide_cutter/peptidecutter.pl). Further research will define the putative role of caspase-1 in this setting. The MA191-induced depletion of FLT3-ITD is not a consequence but occurs prior to apoptosis.

Like in our previous study [29], we found that BIM was necessary for the depletion of FLT3-ITD by PROTACs. The concomitant decrease of FLT3-ITD and cell vitality in the presence of MA49 did not rule out that this requirement of BIM stems from its pro-apoptotic activities at mitochondria. The fast elimination of FLT3-ITD by MA191 discloses that BIM has functions on protein degradation that extend beyond its pro-apoptotic functions. BIM is one of the co-chaperones of HSP70 [55, 56] and HSP70 is necessary for the folding of VHL [37, 57]. It seems plausible that an abruption of HSP70 function can compromise VHL folding. Studies are underway to determine if HSP70 is necessary for the molecular mechanisms of action of MA191 and other FLT3 PROTACs.

Disabling resistance mechanisms and rational combination therapies involving FLT3 PROTACs may improve AML therapy. Since FLT3-ITD is an inducer of both DNA replication stress and DNA repair [32, 58], its elimination probably causes DNA replication stress/DNA damage and a loss of DNA repair functions. Thus, FLT3 PROTACs may combine favorably with DNA-damaging chemotherapies being a mainstay of AML therapy [59]. As FLT3-ITD is only a target in cells carrying this oncogene, non-selective DNA replication stress induction by MA191 in cells without hyperactive FLT3 is unlikely.

Taken together, this work offers a novel FLT3 PROTAC that overcomes cell autonomous and extrinsic resistance mechanisms and reveals its mode of action. **Table 3** summarizes key advantages of MA191.

**Table 3.** Summary of MA49, MA191, and MA68 activity.

|  | MA49 | MA191 | MA68 |
| --- | --- | --- | --- |
| Rapid degradation of FLT3-ITD | - | + | - |
| Toxic to normal blood cells | - | - | + |
| Re-activation of phospho-ERK | + | - | + |
| Killing of FLT3-ITD- and FLT3-ITD/TKD-dependent AML cells | - | + | - |

## Acknowledgments

We thank Prof. Dr. Oliver Ottmann for his great support and excellent discussions throughout this project. Christina Brachetti and Andrea Piée-Staffa provided eminent technical support. This work is made possible with funding from the German Research Foundation/Deutsche Forschungsgemeinschaft (DFG) grant KR2291/15-1, DFG-project number 495271833, and the Brigitte und Dr. Konstanze Wegener-Stiftung (Projekt 110). Work done in the group of O.H.K. is additionally funded by the DFG grant KR2291/12-1, DFG-project number 445785155; KR2291/9-1, DFG-project number 427404172; KR2291/14-1, DFG-project number 469954457; KR2291/16-1, DFG-project number 496927074, KR2291/17-1, DFG-project number 502534123; DFG-project number 393547839 - SFB 1361, sub-project 11; the Walter Schulz Stiftung; the José Carreras Leukemia Foundation; and the H.W. & J. Hector Stiftung. Work done in the group of W.S. is funded by the DFG grant SI 868/26-1 and 468534282, DFG-project number 49527183. M.A. appreciates the support of DAAD and the Ministry of Higher Education and Scientific Research (Egypt) through the GERLS scholarship.

## Author contributions

M.H., W.S., and O.H.K. drafted the manuscript and all authors contributed to its finalization. M.H., M.L., C.H., Y.Z. and S.N. carried out experiments. M.A. and M.S. synthesized and characterized newly synthesized compounds. J.Z, M.L, and C.A carried out primary AML ex vivo characterization. J.Z., M.B., W.S., I.O., and O.H.K. supervised experimental studies. M.B., I.O., and W.B. contributed unique materials. W.S. supervised the synthesis and in vitro characterization of the compounds. M.H., M.A., C.H., and Y.Z. drew figures. W.S. and O.H.K. designed the project outline and acquired its main funding.

## Competing interests

O.H.K. declares the patents WO2019/034538, WO2016020369A1, and WO/2004/027418, and paid advisory work for the BASF Ludwigshafen, Germany. WO2019/034538 (Synthesis, Pharmacology and use of New and Selective FMS-like tyrosine kinase 3 (FLT3) FLT3 Inhibitors) covers substance classes that are discussed in this work. The substances that are covered in these patents are not the same that are shown in the submitted manuscript. The BASF has not influenced our study, and its products are not discussed in the manuscript. Thus, there are no direct competing interests. All other authors declare that they have no conflict of interest.

## Ethics

Peripheral blood and bone marrow samples were obtained from AML patients with written informed consent in accordance with the Declaration of Helsinki under ethical approval REC: 17/LO/1566. Institutional Review Board Statement Zebrafish husbandry (permit number 35-9185.64/BH Hassel) and experiments (permit number 35-9185.81/G-126/15) were performed according to local animal welfare standards (Tierschutzgesetz §11, Abs. 1, No. 1) and in accordance with European Union animal welfare guidelines (EU Directive 2010/63/EU). All applicable national and institutional guidelines for the care and use of zebrafish were followed. C57BL/6 mice were bred and maintained in the Central Animal Facility of the Johannes Gutenberg-University Mainz under specific pathogen-free conditions on a standard diet according to the guidelines of the regional animal care committee. The “Principles of Laboratory Animal Care” (NIH publication no. 85-23, revised 1985) were followed. Mice at 12 weeks of age were sacrificed for organ retrieval according to § 4(3) TierSchG.

## Additional information

### Supplementary information

The online version contains supplementary material available at.

### Data availability

All data are available within the main text and the Supplementary Information files. Original immunoblot files are provided, and all data are available upon scientific reasonable request.

---

**Figure S1.**
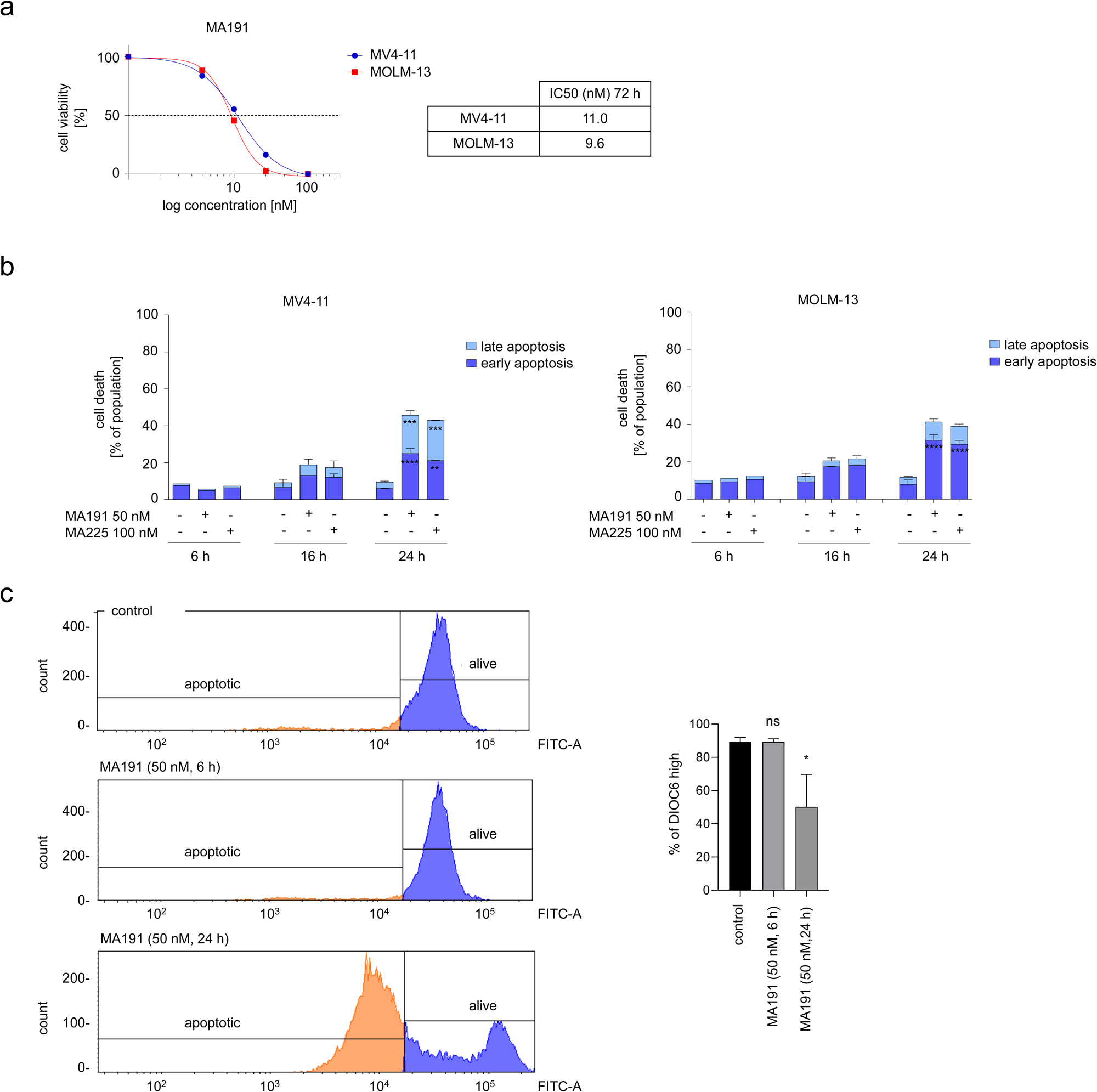
Identification of novel MA191 FLT3 PROTAC. **a** IC_50_ values for apoptosis induction by MA191 in MV4-11 and MOLM-13 cells were determined by annexin-V and PI staining and cytometry. The cells were treated with 0, 5, 10, 20, 50, or 100 nM MA191 for 72 h. IC50 values were calculated with GraphPad Prism 6. **b** MV4-11 and MOLM-13 cells were treated with nM MA191 or 100 nM MA225 for 6, 16, or 24 h and analyzed for apoptosis by annexin-V and PI staining using flow cytometry. The data are representative for the outcome of two ependent experiments; -, untreated, +, treated. Two-way ANOVA statistical test was used to determine statistical significance; *p, <0.05; **p, <0.01; ***p, <0.001; ****p, <0.0001. **c** Representative histograms of DiOC_6_ staining of MV4-11 cells (left) and percentage of the DiOC_6_ high population of MV4-11 and MOLM13 cells (right) treated with 50 nM MA191 for 6 and 24 h respectively. Analysis was made using flow cytometry. The data are representative for the outcome of three independent experiments; One-way ANOVA statistical test was used to determine istical significance; *p, <0.05; **p, <0.01; ***p, <0.001; ****p, <0.0001.

**Figure S2.**
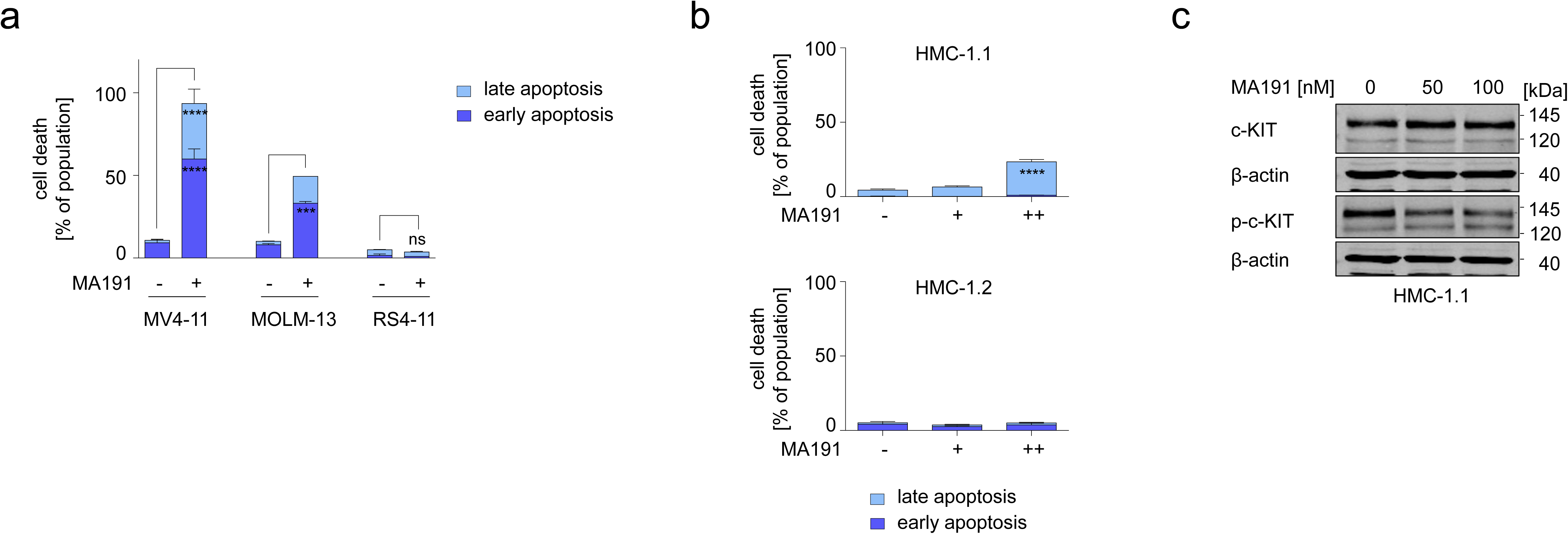
MA191 is most toxic for cells carrying FLT3-ITD. **a** MV4-11, MOLM-13, and RS4-11 cells were treated with 50 nM MA191 for 72 h and analyzed for apoptosis by annexin-V and PI staining using flow cytometry. The data are representative for the outcome of two independent experiments; -, untreated, +, treated. Two-way ANOVA statistical test was used to determine statistical significance; ns, not significant; *p, <0.05; **p, <0.01; ***p, <0.001; ****p, <0.0001. **b** HMC-1.1 (carrying c-KIT with the activating V560G KIT mutation; upper panel) and HMC-1.2 (carrying c-KIT with the activating mutations G560V and D816V; lower panel), were treated with 50 nM or 100 nM MA191 for 72 h and analyzed for apoptosis by annexin-V and PI staining using flow cytometry. The data are representative for the outcome of two independent experiments; -, untreated, +, 50 nM, ++ 100 nM. Two-way ANOVA statistical test was used to determine statistical significance; ****p, <0.0001. **c** HMC-1.1 cells were treated with 50 or 100 nM MA191 for 24 h and analyzed by immunoblot for the expression of c-KIT and p-Y719-c-KIT. The protein levels of β-actin were determined to verify equal loading of samples on all tested membranes. The data are representative for the outcome of three independent experiments; p-, phosphorylated; kDa, molecular weight in kilodalton.

**Figure S3.**
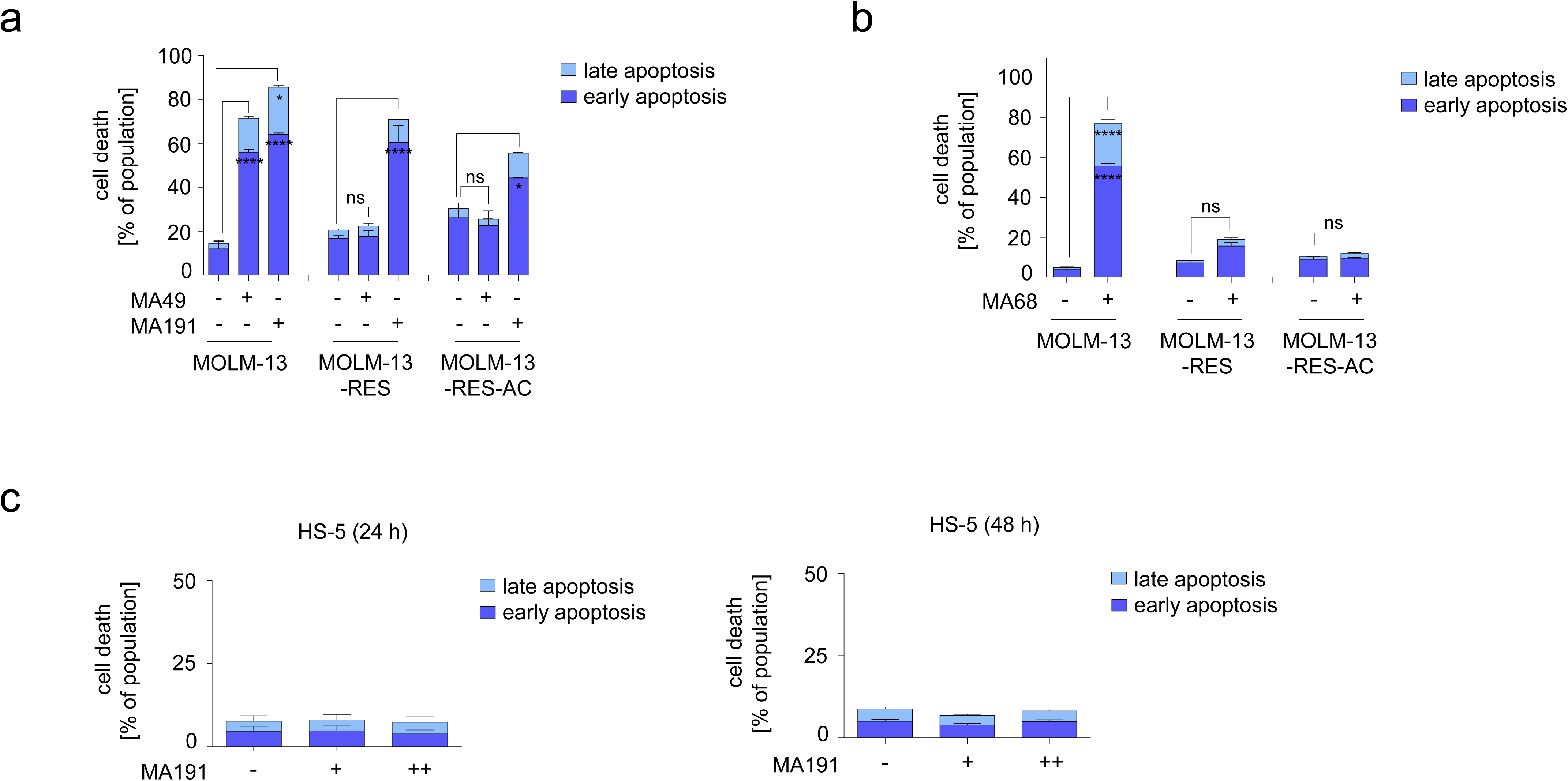
MA191 is effective against FLT3-ITD with secondary D835Y TKD mutation. **a, b** MOLM-13, MOLM-13-RES, and MOLM-13-RES-AC were treated with 200 nM **a** MA49, MA191, or **b** MA68 for 48 h and analyzed for apoptosis by annexin-V and PI staining using flow cytometry. The data are representative for the outcome of two independent experiments; -, untreated, +, 200 nM. Two-way ANOVA test was used to determine statistical significance; ns, not significant; *p, <0.05; **p, <0.01; ***p, <0.001; ****p, <0.0001. **c** HS-5 cells that were co-cultured with MV4-11 cells were treated with 50 or 100 nM MA191 for **c** 24 h or **d** 48 h and analyzed for the apoptosis by annexin-V and PI staining using flow cytometry. The data are representative for the outcome of three independent experiments; -, untreated, +, 50 nM, ++, 100 nM. Two-way ANOVA statistical test was used to determine statistical significance.

